# 3D Spheroid Primary Human Hepatocytes for Prediction of Cytochrome P450 and Drug Transporter Induction

**DOI:** 10.1101/2022.10.30.514199

**Authors:** Erkka Järvinen, Helen S. Hammer, Oliver Pötz, Magnus Ingelman-Sundberg, Tore B. Stage

**Affiliations:** Clinical Pharmacology, Pharmacy and Environmental Medicine, Department of Public Health, University of Southern Denmark, Odense, Denmark; SIGNATOPE GmbH, Reutlingen, Germany; Department of Physiology and Pharmacology, Karolinska Institutet, Stockholm, Sweden; Department of Clinical Pharmacology, Odense University Hospital, Odense, Denmark

**Author notes:** **CORRESPONDING AUTHOR:** Tore Bjerregaard Stage, J.B. Winsløws Vej 19, 2nd floor, DK-5000 Odense C, Denmark. **FUNDING INFORMATION**, This work was funded by Novo Nordisk Foundation (Grant NNF19OC0058275), Lundbeck Foundation Fellowship (Grant R307-2018-2980), the ERC Advanced Grant (AdG) project HEPASPHER (grant agreement 742020) and by The Swedish Research Council, grant 2018-05766.

**Keywords:** Primary human hepatocytes, induction, DDI, CYP, transporter, solute carrier transporters, efflux transporters, uptake transporters, CES1

## Abstract

Primary human hepatocytes (PHHs) have been the gold standard *in vitro* model for the human liver and are crucial to predict hepatic drug-drug interactions. The aim of this work was to assess the utility of 3D spheroid PHHs to study induction of important cytochrome P450 (CYP) enzymes and drug transporters. 3D spheroid PHHs from three different donors were treated for four days with rifampicin, dicloxacillin, flucloxacillin, phenobarbital, carbamazepine, efavirenz, omeprazole or β-naphthoflavone. Induction of CYPs 1A1, 1A2, 2B6, 2C8, 2C9, 2C19, 2D6, 3A4, P-gp/*ABCB1*, MRP2/*ABCC2, ABCG2*, OCT1/*SLC22A1, SLC22A7, SLCO1B1* and *SLCO1B3* were evaluated at mRNA and protein levels. Enzyme activity of CYP3A4, CYP2B6, CYP2C19 and CYP2D6 were also assessed. Induction of CYP3A4 protein and mRNA correlated well for all donors and compounds and had a maximal induction of 5-to 6-fold for rifampicin, which closely correlates to induction observed in clinical studies. Similar estimates were found for dicloxacillin and flucloxacillin, which also correlates to findings from clinical studies. Rifampicin induced the mRNA of *CYP2B6* and *CYP2C8* by 9- and 12-fold, while the protein levels of these CYPs reached 2- and 3-fold induction, respectively. Rifampicin induced CYP2C9 protein by 1.4-fold, while the induction of CYP2C9 mRNA was over 2-fold in all donors. Rifampicin induced *ABCB1*, *ABCC2* and *ABCG2* by 2-fold. In conclusion, 3D spheroid PHHs is a valid model to investigate mRNA and protein induction of hepatic drug metabolizing enzymes and transporters, and this model provides a solid basis to study induction of CYPs and transporters, which translates to clinical relevance.

## INTRODUCTION

Drug-drug interactions (DDIs) are typically caused by inhibition or induction of drug metabolizing enzymes and transporters.^1^ Strong cytochrome P450 enzyme (CYP) inducers may reduce drug exposure of victim drugs dramatically by over 10-fold.^2,3^ Typically, induction of CYP3A4, CYP2B6 and CYP1A2 by investigational compounds are evaluated *in vitro* to evaluate risk of clinical DDIs.^4^ Evaluation of these three enzymes is based on selectivity of different nuclear receptors that mediate drug induction - pregnane X receptor (PXR), constitutive androstane receptor (CAR) and aryl hydrocarbon receptor (AhR) for CYP3A4, CYP2B6 and CYP1A2, respetively.^1^ Nevertheless, other drug-metabolizing enzymes and transporters, such as CYP2Cs and P-glycoprotein (P-gp), are also inducible and prone to clinical DDIs.^1,5^

*In vitro* models of the human liver are crucial for prediction of hepatic drug metabolism and DDIs. Hepatic *in vitro* models are particularly important to assess drug induced enzyme and transporter regulation. Currently, the gold standard models to study the human liver *in vitro* are cultures of primary human hepatocytes (PHHs) isolated from human livers.^6^ Despite progress over the years, satisfactory prediction of induction DDIs and their magnitude remains elusive.^2,3,7–9^ Particularly, studies in 2D monolayer PHHs may result in highly variable induction or downregulation responses that may not represent the clinical magnitude and variability of DDIs.^3,8–11^ For example, dicloxacillin induced CYP3A4 and CYP2C9 by 30- and 2-fold in 2D monolayer PHHs, while dicloxacillin decreased the AUC of midazolam and tolbutamide by 1.9- and an 1.4-fold in a clinical study.^12^

PHHs are cultured in different formats, typically as 2D monolayers.^6,13^ However, PHHs lose their phenotype within days in most culture formats.^14–19^ To improve *in vitro* modelling of human liver, a 3D spheroid culture of primary human hepatocytes (3D spheroid PHHs), without addition of extracellular matrices or scaffolds, has been characterized and investigated for several pharmacological applications.^16,18–26^ 3D spheroid PHHs retain the hepatic phenotype of native tissue at the protein and RNA levels, and are viable for several weeks.^16,24^ 3D spheroid PHHs have outperformed 2D monolayer and sandwich cultured PHHs in hepatotoxicity assays^15,20^ and they retain drug metabolism activity of important CYPs for several weeks allowing long-term experiments.^15,16,18,19,24,25^ Assessment of drug clearance and identification of low-abundant metabolites is also feasible in 3D spheroid PHHs.^22,25,26^ In contrast to 2D monolayer PHHs, 3D spheroid PHHs express *CYP3A4* mRNA at almost 200-fold higher levels.^23^ Remarkably, 3D spheroid PHHs identified induction of CYP3A4 by drugs that did not trigger a clear response in 2D monolayer culture of PHHs.^23^

Here, our aim was to characterize comprehensively drug-mediated induction of clinically important CYP enzymes and transporters in 3D spheroid PHHs. To this end, we included several drugs known to cause clinically relevant DDIs. The magnitude of induction was evaluated at the levels of mRNA and protein for CYPs 1A2, 2B6, 2C8, 2C9, 2C19, 2D6 and 3A4 and P-glycoprotein (P-gp) and multidrug resistance-associated protein 2 (MPR2). CYP enzyme activity of 2B6, 2C19, 2D6 and 3A4 was also measured after induction by employing the Basel cocktail.

## METHODS

### 3D SPHEROID CULTURE OF PRIMARY HUMAN HEPATOCYTES

Three lots, HU8345-A, HU8339-A and Hu8373-A of cryopreserved PHHs qualified for spheroid formation (HMCPSQ) were acquired from Thermo Fisher Scientific (Walthan, MA, USA). All cell culture reagents were also from Thermo Fisher Scientific. For the spheroid cultures, 1,500 cells were seeded on day 0 and induction studies were conducted between days 8 and 12. The spheroid culture protocol was adapted from previous studies.^16,23^ The detailed description of 3D spheroid culture of PHHs and donor information (**Table S1)** is in **Supplementary materials**.

### INDUCTION TREATMENTS OF 3D SPHEROID PRIMARY HUMAN HEPATOCYTES

Spheroids were treated for 96 h with different inducers. All compounds (Sigma-Aldrich, St. Louis, MO, USA) were prepared as 1000x stocks in dimethyl sulfoxide (DMSO, from Sigma-Aldrich) and the final concentration of DMSO was 0.1% in each experiment. The final concentrations of inducers were 10 μM rifampicin, 100 μM dicloxacillin, 100 μM flucloxacillin, 250 μM phenobarbital, 50 μM carbamazepine, 10 μM efavirenz, 50 μM omeprazole and 10 μM β-naphthoflavone. Experiments also included vehicle control samples (0.1% DMSO) and no treatment controls. After 48 hours on day 10, medium was changed to all treatments. For RNA isolation and protein quantification, triplicate samples of pools of 16 spheroids were collected on day 12 from all 3 donors. RNA isolation samples were dissociated with 500 μl of Qiazol reagent (Qiagen, Germantown, MD, USA) after complete removal of cell culture media. For protein quantification, cell culture media was removed, and spheroids were washed once with 1 ml of warm phosphate-buffered saline solution containing magnesium and calcium (Thermo Fisher Scientific). All samples were stored at −80°C before RNA or protein extraction.

### CYP ENZYME ACTIVITY EVALUATION

Enzyme activity of CYP1A2, CYP2B6, CYP2C9, CYP2C19, CYP2D6 and CYP3A4 was determined with the Basel cocktail in spheroids from donor 1 and 3 on day 12.^27^ Losartan, metoprolol and caffeine (Sigma-Aldrich) were dissolved in water as 6,000x or 500x (caffeine) stocks. Omeprazole and efavirenz (Sigma-Aldrich) and midazolam (Toronto Research Chemicals, Toronto, ON, Canada) were dissolved in DMSO as 6,000x stocks. The final concentration of DMSO was 0.15% in enzyme activity assays. Before applying the cocktail to spheroids, cells were washed three times with 100 μl of the maintenance medium. The residual volume was 20 μl between the washes and before the third wash spheroids were incubated for 2 to 3 hours in a cell culture incubator. Final concentrations of the Basel cocktail, in a volume of 100 μl, were 160 μM caffeine, 20 μM efavirenz, 30 μM losartan, 30 μM omeprazole, 40 μM metoprolol and 10 μM midazolam (modified from ^27^). Triplicate samples, each containing medium and spheroid from a well, were collected after 0.15-, 0.5-, 8- and 24-hour incubations with the Basel cocktail. The samples were stored at −80°C before analysis. Analytical methods for metabolite analysis are described in **Supplementary methods**.

### SPHEROID VIABILITY AND MORPHOLOGY

ATP content of spheroids was used as a surrogate for viability.^16,21^ Single spheroids were transferred to wells of an opaque, white plate containing 25 μl of the maintenance medium per well. A standard curve for ATP (Thermo Fisher Scientific) was prepared in the maintenance medium. CellTiter-Glo 3D cell viability assay from Promega (Madison, WI, USA) was added as 25 μl aliquots to each well and the plate was vortexed for 5 mins. After half an hour, luminescence of each well was measured and absolute ATP content per well was calculated based on the ATP standard curve.

Morphology of spheroids was observed every second or third day by imaging spheroids with a 10x magnification of a light microscope equipped with a digital camera from Motic (Hong Kong, China). Brightness and contrast adjustments were equal for every picture.

### RNA EXTRACTION AND qPCR

RNA was extracted with the phenol-chloroform method according to the manufacturer’s protocol (Qiagen) with small modifications. Total volume of Qiazol reagent was 1 ml and RNA was co-precipitated with 15 μg of RNA grade glycogen (Thermo Fisher Scientific). In addition, RNA pellet was washed three times with 75% ethanol and heating was used to evaporate remaining ethanol as well as to aid solubilization of the pellet in water.^28^ cDNA was synthesized from ~400 ng of RNA with High-Capacity cDNA Reverse Transcription Kit with RNase Inhibitor (Thermo Fisher Scientific). For qPCR reactions, 10 ng of cDNA (based on RNA concentrations) was transferred to each well containing TaqMan Universal Master Mix II (Thermo Fisher Scientific) and a target specific TaqMan assay (assays reported in **Supplementary methods**) in a volume of 10 μl. qPCR reactions were run according to the manufacturer’s protocol (Thermo Fisher Scientific) for 40 cycles, and each plate included glyceraldehyde-3-Phosphate dehydrogenase (GAPDH) amplification for every sample, and no-template and no-reserve transcriptase controls for every target.

For the data analysis, Ct-values of GAPDH were subtracted from Ct values of targets for each sample resulting in deltaCt values. The deltaCt values were transformed by 2^-Ct^ and all samples were normalized to the mean value of control samples (0.1% DMSO, Day 0 or no treatment samples) separately for each target.

### PROTEIN QUANTIFICATION BY LIQUID CHROMATOGRAPHY-MASS SPECTROMETRY

A targeted LC-MS method for human CYP and transporter protein quantification, including selective immunoprecipitation of peptides, was previously reported.^29^ Same number of spheroids and cells were used for every analysis. Briefly, spheroids or cells were denatured in 42 mM ammonium bicarbonate buffer containing 1.14 mM dithiothreitol, and 9 mM iodoacetamide for 15 min at 90°C (adopted from ^30^). Subsequently, samples were digested with trypsin (MS-grade Pierce Trypsin Protease from Thermo Fisher Scientific) for 16 h at 37°C. Reactions were stopped by addition of phenylmethanesulfonyl fluoride to a final concentration of 1 mM and a total volume of 70 μL per sample. Surrogate peptides and internal standard peptides were precipitated using triple X proteomics antibodies from 25 μL of digested samples.^29^ Peptides were then subjected to LC-MS analysis and quantification. The surrogate peptides employed for protein quantification and LC-gradients are presented in **Table S3**. Raw data were processed using Skyline software (MACOSS Lab, Department of Genome Sciences, University of Washington, Seattle, USA) and TraceFinder 4.1 (Thermo Fisher Scientific).

## RESULTS

### 3D SPHEROID CULTURE OF PRIMARY HUMAN HEPATOCYTES

On day 7 and 9, spheroids showed tightly packed morphology with a diameter of 200 μm or higher **(Figure 1)**. ATP content of spheroids was measured as a surrogate for cell viability **(Figure 2)**. ATP levels dropped slightly within time for donors 1 and 3 **(Figure 2)**, which aligns with shrinking of these spheroids from day 7 to 16 **(Figure 1)**. ATP content of donor 2 was stable for all measured culture days **(Figure 2)**, and spheroids from this donor did not shrink clearly **(Figure 1)**. ATP content of spheroids from donor 2 and 3 were followed until day 19 and 35, respectively **(Figure S1)**.

**FIGURE 1.**
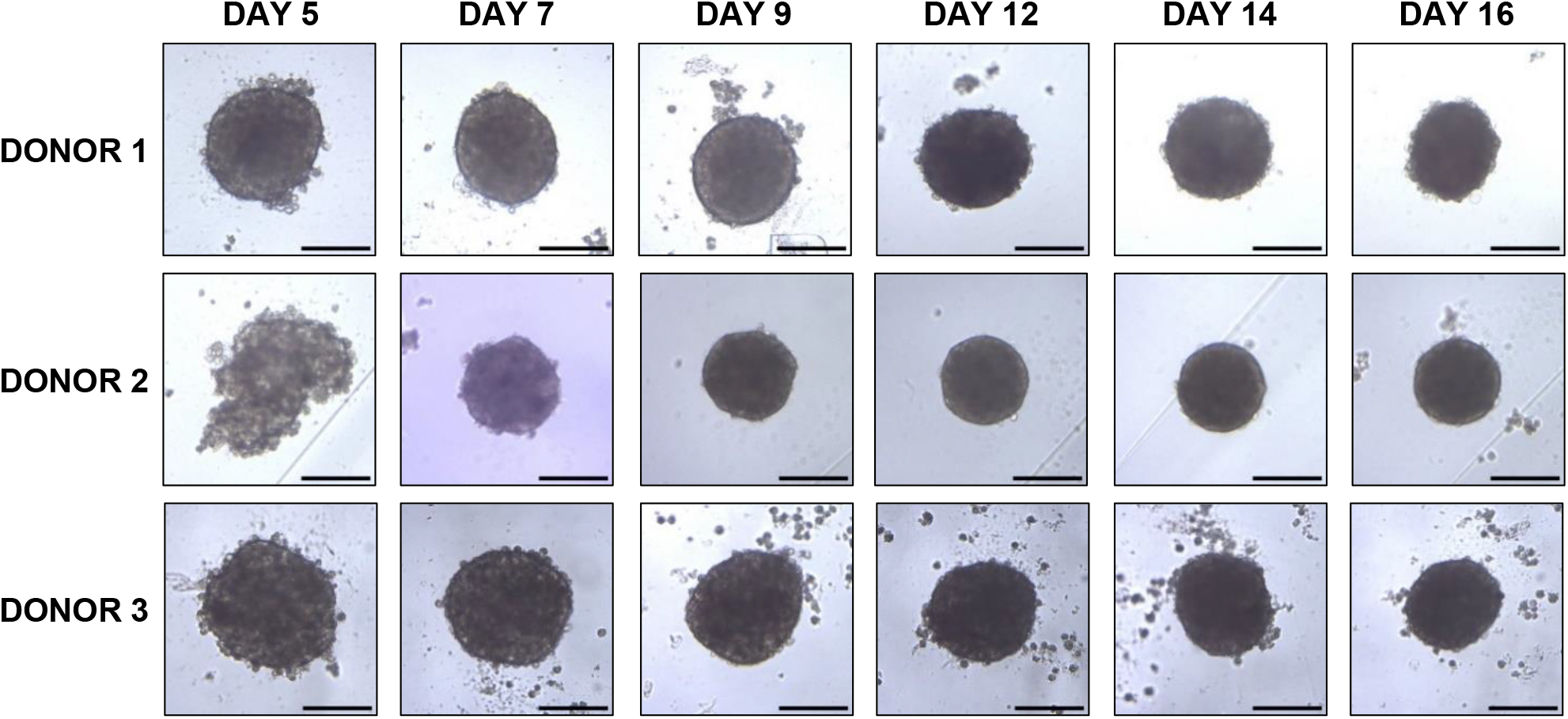
Morphology of 3D spheroid PHHs from day 5 to day 16. Primary human hepatocytes were seeded on day 0 and spheroids formed within five days. Bright field microscopy pictures of spheroids from each donor were taken. For each donor, the pictures are from a same well, except the day 16 picture of donor 1 is from a different well. Scale bars present 200 μm.

**FIGURE 2.**
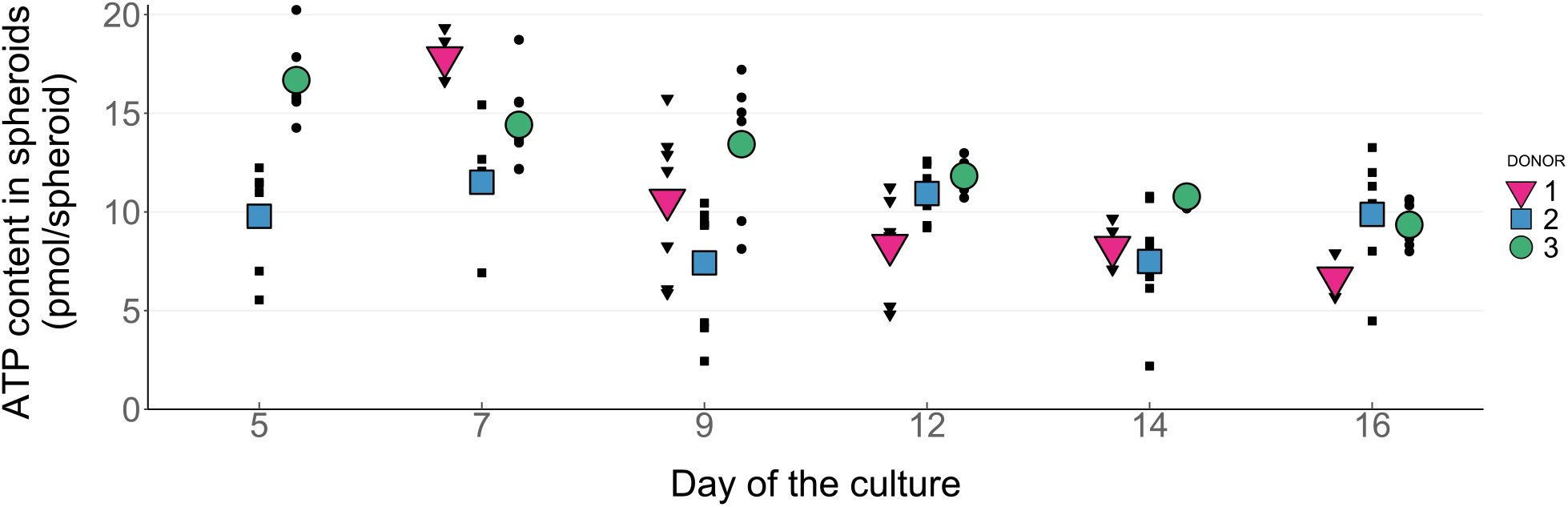
Viability of 3D spheroid PHHs from day 5 to 16. ATP content of spheroids was determined for each donor on days 5, 7, 9, 12, 14 and 16 after seeding hepatocytes in ultra-low attachment plates. Data for donor 1 on day 5 is missing. N = 4-8 biological replicates (spheroids) per donor and time-point.

The baseline expression of mRNA and protein levels was evaluated for CYPs and drug transporters from freshly thawed cryopreserved PHHs on day 0 and from 3D spheroid PHHs on day 12 **(Figures S2-S4, Table S4)**. The expression of CYP3A4 was same or higher in spheroids in comparison to freshly thawed cryopreserved PHHs on day 0 both at the mRNA **(Figures S2 and S4)** and protein **(Figure S3)** levels. CYPs 2B6, 2C8 and 2C9 were downregulated in spheroids compared with freshly thawed cryopreserved PHHs on day **(Figures S2-S4).**

### DRUG-MEDIATED INDUCTION of mRNA and PROTEIN LEVELS IN 3D SPHEROID PRIMARY HUMAN HEPATOCYTES

Spheroids were exposed to different inducers that were chosen based on their activation of nuclear receptors. Rifampicin, dicloxacillin and flucloxacillin are strong, medium and weak PXR activators^1,12^ respectively, while phenobarbital is both a PXR and CAR activator.^23^ Carbamazepine and efavirenz activate CAR^1^, while β-naphthoflavone is an AhR activator.^31^ Finally, omeprazole is an inducer of CYP1A2^1^, however, other pathways beyond activation of AhR are likely involved in omeprazole’s effect on CYP enzymes and drug transporters.^31,32^

Spheroids were treated for 96 hours between the culture days 8-12 and treatments included a medium change on day 10. Effect of the treatments on the viability of spheroids was evaluated for each donor **(Figure S5)**. Only β-naphthoflavone decreased the ATP content of spheroids by more than 20% (31% for β-naphthoflavone).

### CYTOCHROME P450 ENZYME INDUCTION

At the mRNA levels, PXR-activating compounds induced *CYP2B6, CYP2C8, CYP2C9, CYP2C19* and *CYP3A4* **(Figure 3A)** and the magnitude of induction was similar to expected DDI potential of these compounds – rifampicin > dicloxacillin > flucloxacillin. *CYP2B6* was highly induced by CAR-activating compounds phenobarbital, carbamazepine and efavirenz but also by rifampicin. Omeprazole also caused substantial induction of *CYP2B6* and *CYP2C19*, while *CYP2C8*, *CYP2C9* and *CYP3A4* were less induced. *CYP1A1* and *CYP1A2* mRNA were induced only by omeprazole and β-naphthoflavone and induction was over 10-fold the vehicle control **(Figure 3A)**. None of the compounds induced *CYP2D6* **(Figure 3A)**. β-naphthoflavone had a different effect in comparison to other inducers as this compound downregulated *CYP3A4* and to a less extent *CYP2C8*, *CYP2C9* and *CYP2D6. CYP2C19* was not inducible for donor 2 **(Figure 3A)**.

**FIGURE 3.**
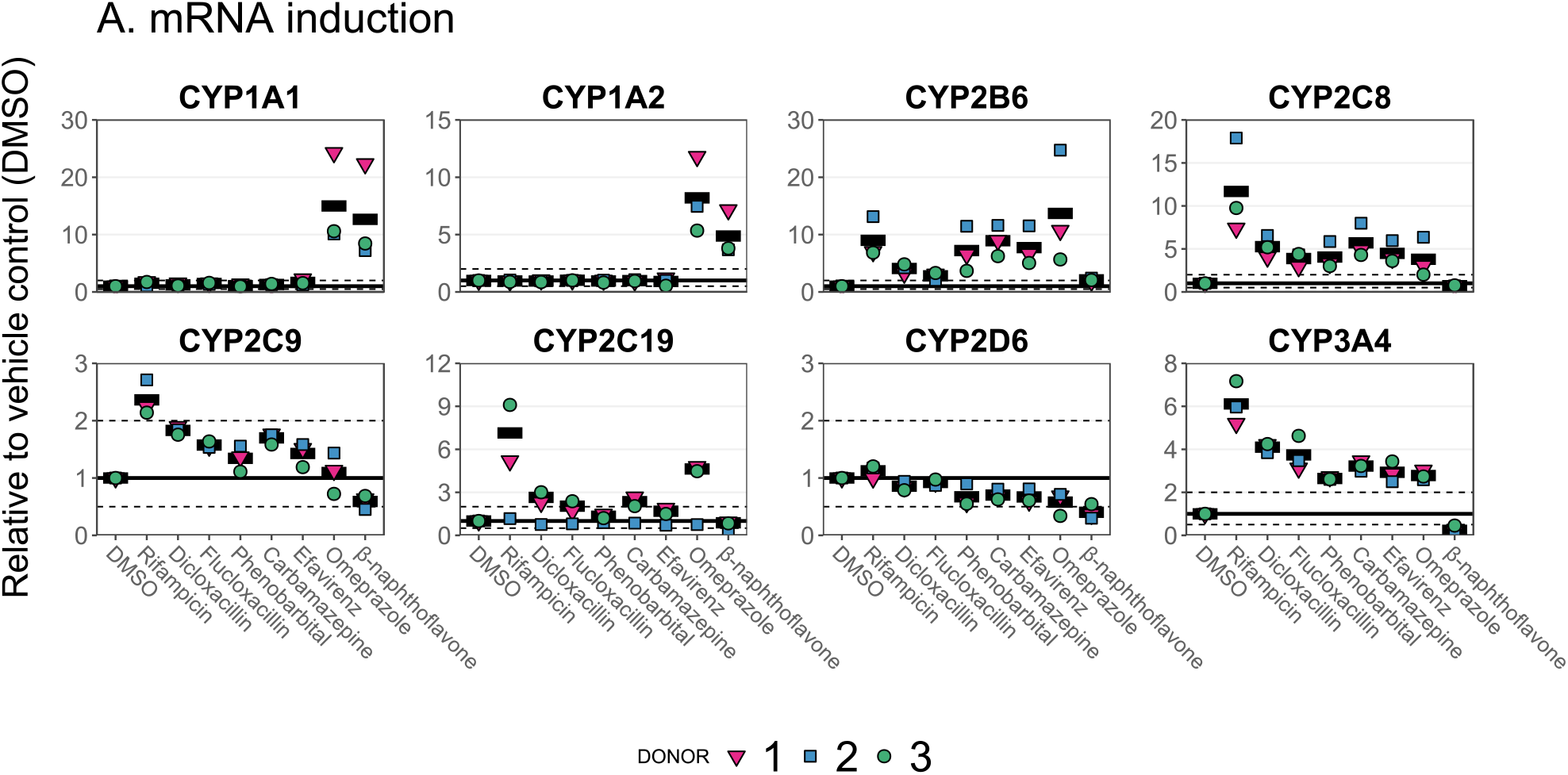
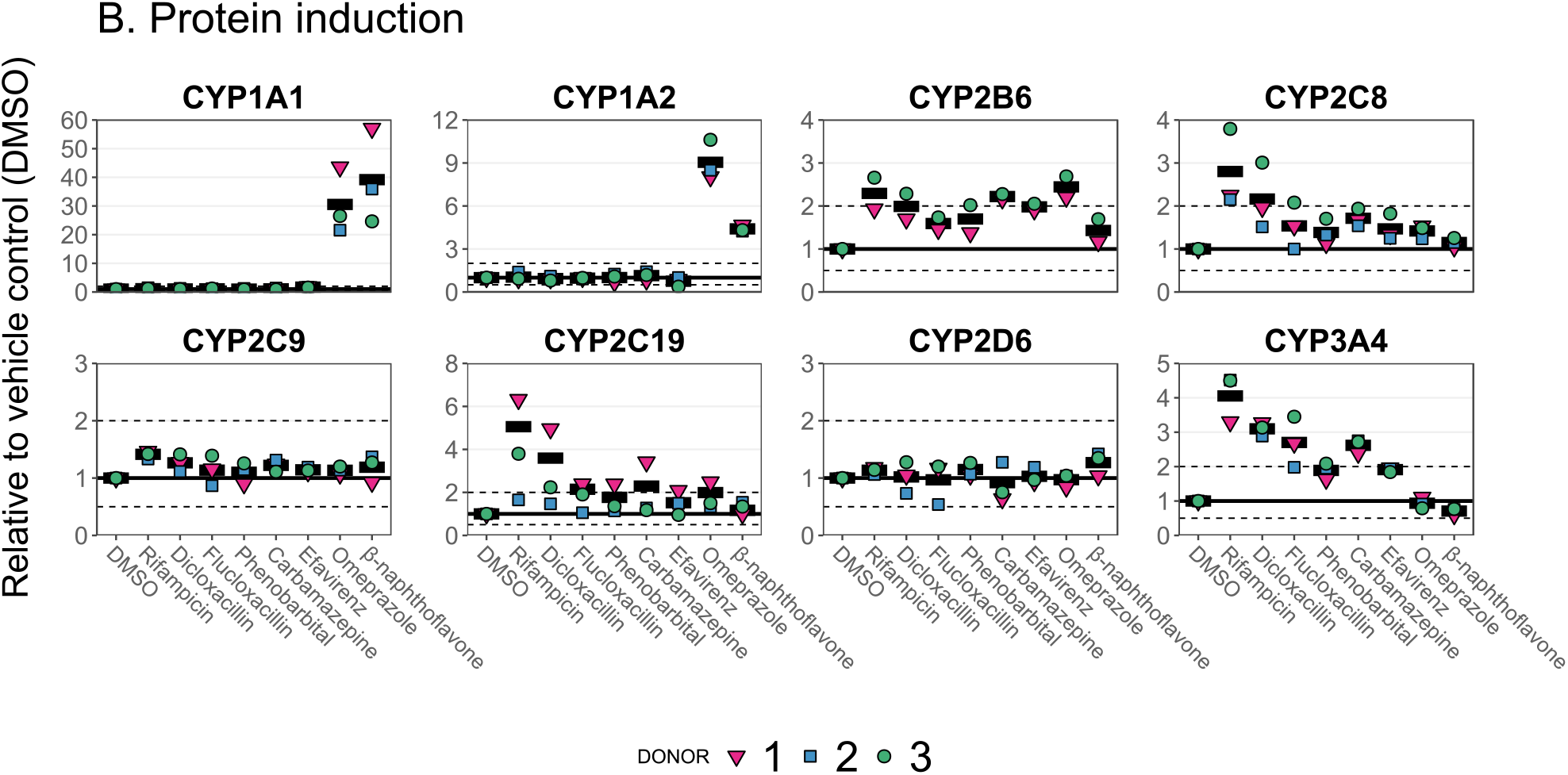
Compound-specific induction of main hepatic CYP enzymes in 3D spheroid PHHs. Induction of mRNA (A) or protein (B) levels were determined after 96 h of treatment with the compounds. For each donor, induction was normalized to the mean response of 0.1% DMSO group. Data are summarized as means of three hepatocyte donors, except for CYP2C19 of donor 2 that was excluded due to no induction. Dashed lines present two-fold difference to the DMSO group. N = 3 hepatocyte donors each presented as a mean value of three biological replicates of pools of 16 spheroids.

Protein levels of CYPs were induced with a similar pattern as mRNA but overall to a lower extent **(Figure 3B)**. CYP1A1 was the only protein that was induced to a higher extent at the protein levels in comparison to mRNA levels **(Figures 3A and 3B)**. CYP2C19 induction was absent in donor 2 in accordance with the mRNA data **(Figures 3A and 3B)**. CYP2B6 levels were below limit of quantification in donor 2. No compound downregulated CYPs at the protein level **(Figure 3B)**. Results on the protein level induction of other drug-metabolizing enzymes (CYP2C18, CYP2E1, CYP3A5, CYP3A7, cytochrome P450 reductase (POR) and carboxylesterase 2 (CES2)) are presented in **Figure S6**.

The effect of vehicle used in induction studies on the expression of CYP enzymes in 3D spheroid PHHs was evaluated by comparing the expression between non-treated and 0.1% DMSO-treated spheroids **(Figure S7)**. The mRNA expression of *CYP2B6* was highly induced by 0.1% DMSO in donors 1 and 3 (4-fold), while the induction was less than 2-fold in donor 2 **(Figure S7A)**. The mRNA expression of *CYP2C8*, CYP2C9, *CYP3A4* and *CYP2D6* was induced 1.5-2-fold by 0.1% DMSO **(Figure S7A)**. An effect of 0.1% DMSO on the protein levels of CYPs was found for CYP3A4 that was induced 1.4- and 2.8-fold in donors 1 and 3 **(Figure S7B)**.

### DRUG TRANSPORTER and CARBOXYLESTERASE 1 INDUCTION

Efflux transporter *ABCB1, ABCC2* and *ABCG2*, and *CES1* mRNA levels were induced two-fold by rifampicin, while dicloxacillin and flucloxacillin treatments resulted in lower induction **(Figure 4A)**. Uptake transporters were not induced **(Figure 4A)**, however, omeprazole downregulated the mRNA expression of *SLC22A1*, *SLC22A7*, *SLCO1B1* and *SLCO1B3* to varying extent. β-naphthoflavone downregulated all uptake transporters and *ABCB1* at the mRNA levels, while it caused weak induction of *ABCG2* **(Figure 4A)**.

**FIGURE 4.**
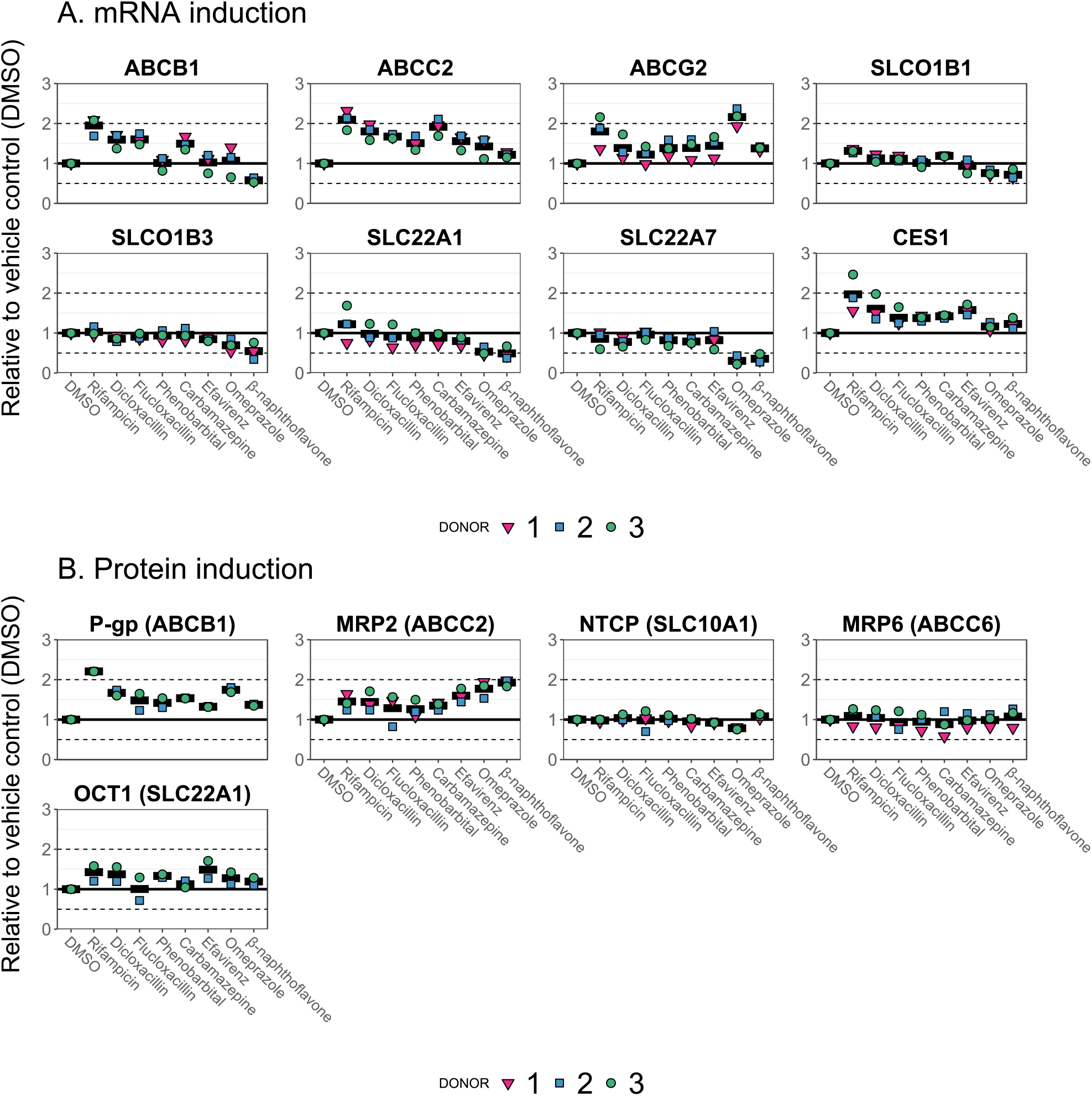
Compound-specific induction of hepatic transporters in 3D spheroid PHHs. Induction of mRNA (A) or protein (B) levels were determined after 96 h of treatment with the compounds. For each donor, induction was normalized to the mean response of 0.1% DMSO group. Data are summarized as means of three hepatocyte donors. P-gp and OCT1 were below limit of quantification in donor 1. Dashed lines present two-fold difference to the DMSO group. N = 3 hepatocyte donors each presented as a mean value of three biological replicates of pools of 16 spheroids.

The protein levels of P-gp were induced similarly by different compounds and to a similar extent **(Figure 4B)** as the mRNA levels **(Figure 4A)**, while the protein levels of MRP2 were induced to a lower extent **(Figure 4B)** than the mRNA levels **(Figure 4A)** by most compounds. The protein expression of organic cation transporter 1 (OCT1) was induced by several compounds to a maximum of 1.5-fold over the control **(Figure 4B)**, although no any induction was found at the mRNA levels **(Figure 4A)**. No compound induced the protein expression of sodium/taurocholate cotransporting polypeptide (NTCP) or MRP6.

The vehicle (0.1% DMSO) did not affect the mRNA or protein expression of transporters or esterases in comparison to non-treated spheroids **(Figure S7)**.

### CORRELATION BETWEEN INDUCTION of mRNA and PROTEIN LEVELS

Correlation between induction of mRNA and protein levels by different inducers was evaluated **(Figure 5)**. Overall, induction of *ABCB1*/P-gp, CYP1A2, CYP3A4 and CYP2C19 showed close correlation between the relative change of mRNA and protein levels **(Figure 5)**. The high induction of mRNA levels of *CYP2C8* and *CYP2B6* did not translate to high levels of protein expression **(Figure 5)**.

**FIGURE 5.**
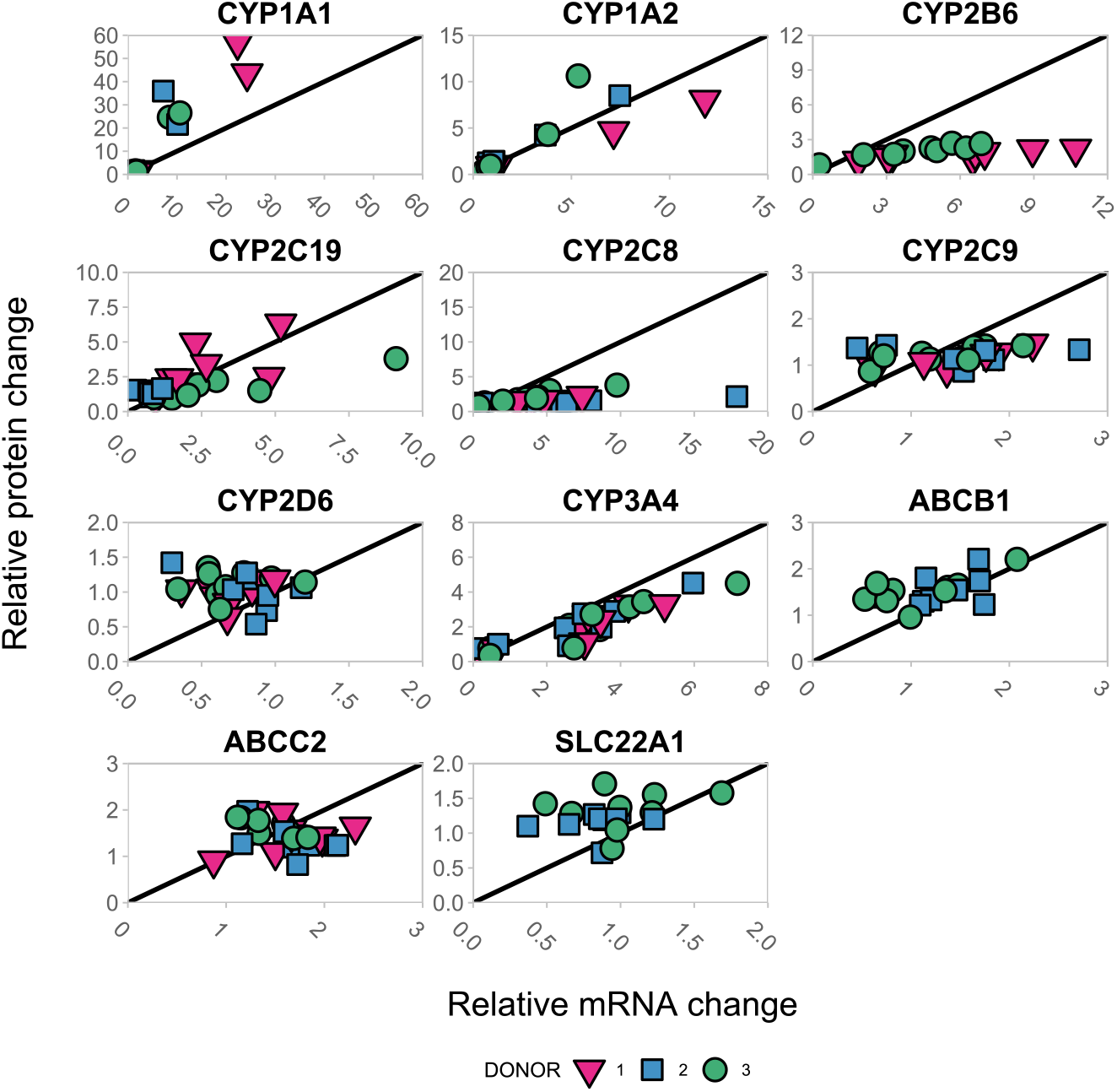
Correlation between the relative mRNA and protein induction in three different donors treated with 8 different drugs. Each data point presents the mean of relative induction at mRNA (x axis) and protein (y axis) level for one of the compounds, excluding 0.1% DMSO, from Figures 3 and 4. Solid lines represent the line of unity.

Linear regression between the relative change in protein and mRNA levels was analyzed for the most highly induced CYPs – 2B6, 2C8, 2C19 and 3A4 **(Figure S8)**. The regression analysis of CYP2B6 resulted in slopes of 0.10 and 0.25 for donor 1 and 3, respectively **(Figure S8)**. The same modelling for CYP2C8 resulted in slopes of 0.19, 0.06 and 0.31 for donor 1, 2 and 3, respectively **(Figure S8)**. In the case of CYP2C19, the slopes were 0.79 and 0.31 for donor 1 and 3, respectively **(Figure S8)**. For CYP3A4, the slopes were 0.58, 0.63 and 0.62 for donor 1,2 and 3, respectively **(Figure S8)**. The slopes indicate that the mRNA levels overestimated by 4- to 10-fold the induction of protein expression for CYP2B6 and by 3- to 17-fold for CYP2C8, while for CYP3A4 the overestimation was less than 2-fold.

### DRUG-MEDIATED INDUCTION OF CYP ENZYME ACTIVITIES IN 3D SPHEROID PRIMARY HUMAN HEPATOCYTES

The Basel cocktail contained probe drugs for CYP1A2, CYP2B6, CYP2D6, CYP2C9, CYP2C19 and CYP3A4. Paraxanthine and losartan carboxylic acid were below limit of quantification in almost every sample and therefore no results are available for CYP1A2 or CYP2C9. Based on the linear fittings of metabolite formation within time **(Figure S9)**, data for the fold analysis **(Figure 6)** were extracted from 0.5-hour incubation for CYP2C19 and from 8-hour incubations for CYP2B6, CYP2D6 and CYP3A4. CYP-catalyzed metabolite formation was confirmed by inhibiting enzyme activity of CYPs with either pan-CYP inhibitor 1-aminobenzotriazole or verapamil **(Figure S10)**.

**FIGURE 6.**
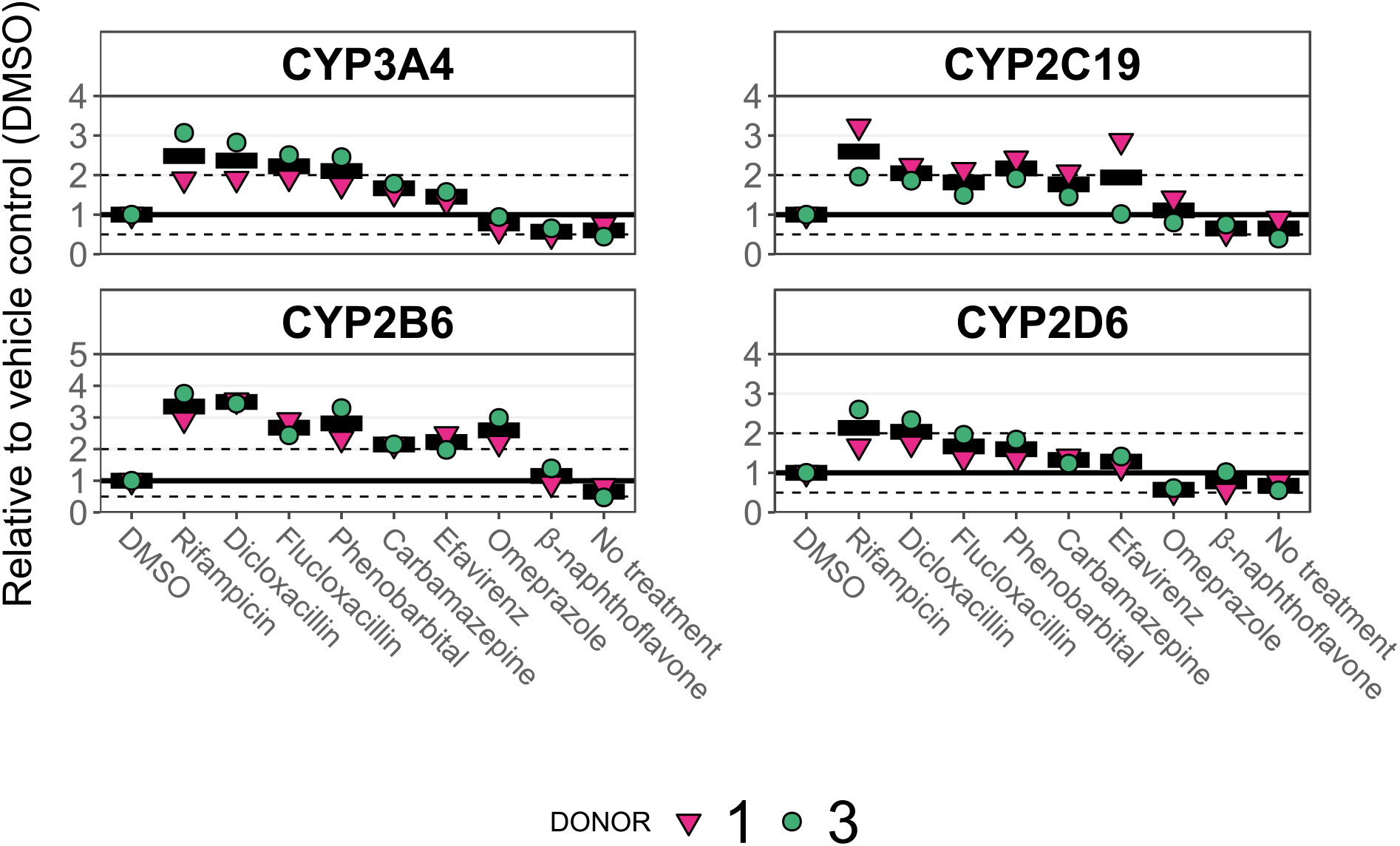
Induction of CYP3A4, CYP2C19, CYP2B6 and CYP2D6 enzyme activities. Spheroids were treated with inducers or vehicle (0.1% DMSO) or were untreated (No treatment) for 96 hours before incubation with the Basel cocktail (10 μM midazolam, 30 μM omeprazole, 20 μM efavirenz, 40 μM metoprolol, 30 μM losartan and 160 μM caffeine). The metabolic activities of CYP3A4, CYP2C19, CYP2B6 and CYP2D6 were measured from samples incubated for 8 hours or 0.5 hour in the case CYP2C19. Data were normalized to mean concentration of metabolites in the vehicle group and are presented as means of two hepatocyte donors. N = 2 hepatocyte donors each presented as a mean value of three biological replicates per condition. Dashed lines indicate a two-fold difference to the vehicle group.

Enzyme activities of CYP2C19 and CYP3A4 were induced **(Figure 6)** similarly to the protein levels **(Figure 3B)** by different compounds but the maximal inductions were higher at the protein levels (**Figure S11**). Induction of CYP2B6 activity was more pronounced **(Figure 6)** than the induction of protein levels **(Figure S11)**. The enzyme activity of CYP2D6 was induced up to 2-fold, although no protein induction was found **(Figures 3B and S11)**, and the magnitude of induction was similar to the induced activity of CYP3A4 **(Figure 6)**. Enzyme activities in spheroids without any treatment were 1.5- to 1.7-fold lower than in vehicle (0.1% DMSO) treated spheroids **(Figure 6)**.

## DISCUSSION

We investigated the utility of 3D spheroid PHHs to study regulation of drug metabolizing CYPs and drug transporters. We evaluated the induction of CYPs by different inducers at the level of mRNA, protein, and enzyme activity and found similar induction pattern in all endpoints but with varying extent of induction. CYP3A4 was most consistently induced for all endpoints, while CYP2C8 and CYP2B6 had discrepancy between mRNA and protein induction. We also evaluated the induction of several uptake and efflux transporters and found reproducible response between three donors and no evidence of uptake transporter induction.

CYP3A4 is the most important enzyme in drug metabolism and thus prediction of induction mediated DDIs for CYP3A4 is highly important.^1,4^ We found that the maximal induction of CYP3A4 mRNA and protein by rifampicin were 6- and 4-fold **(Figure 3)**, which agrees with a study that found a 4-fold induction of CYP3A4 protein in hepatic extracellular vesicles isolated from plasma of rifampicin treated subjects.^33^ Moreover, dicloxacillin was identified as a weak CYP3A4 inducer in a clinical study although results from 2D monolayer PHHs showed up to 30-fold induction of *CYP3A4* mRNA.^12^ Here, dicloxacillin induced both mRNA and protein levels of CYP3A4 weaker than rifampicin **(Figure 3)**, which reflects the clinical interaction more accurately than the magnitude of induction observed in 2D monolayer PHHs. This highlights that 3D spheroid PHHs are able to distinguish between strong and weak inducers.

3D spheroid PHHs retain the *in vivo* hepatic phenotype of human liver in culture for several weeks and stably express CYPs and transporters, which is a prerequisite for *in vitro* to *in vivo* extrapolation.^15,16,18,19,24^ Previously, it was shown that high baseline expression of *CYP3A4* mRNA in 3D spheroid PHHs resulted in clinically more relevant magnitude of induction than in 2D monolayer PHHs.^23^ Moreover, 3D spheroid PHHs were able to identify inducers that failed to induce CYP3A4 in 2D monolayer PHHs.^23^ We confirmed that 3D spheroid PHHs express CYP3A4 protein after 12 days of culture comparable to that of freshly thawed PHHs **(Figures S3)**.

The mRNA of *CYP2B6* and *CYP2C8* were highly induced by different compounds, while the protein levels were induced to a much lower extent **(Figure 3)**. The expression of CYP2C8 and CYP2B6 were downregulated in spheroids on day 12 compared with freshly thawed PHHs on day 0 similarly to previous reports from spheroid and other advanced hepatic cultures **(Table S4)**. This downregulation may explain the high extent of induction of *CYP2B6* and *CYP2C8* mRNA by different inducers. In 2D monolayer PHHs, up to 23- to 40-fold induction of *CYP2B6* mRNA by rifampicin, carbamazepine and efavirenz has been reported^7^ compared with the maximal induction of 11- to 13-fold by same compounds here **(Figure 3A)**. Similarly, omeprazole, phenobarbital and rifampicin induced highly CYP2B6 protein levels in sandwich cultured PHHs.^34^ However, the extent of observed pharmacokinetic changes mediated by CYP2B6 induction are much lower than 10-fold^7^, which highlights that the quantification of protein induction or enzyme activity may be a better surrogate for predictions of clinical DDIs than mRNA induction. Our findings support the weakness of mRNA induction for *in vitro* to *in vivo* translation, since only the protein level induction of CYP2B6 was closer to the magnitude of induction of enzyme activity **(Figures 3 and 6)**.

CYP2C9 was only weakly induced by the compounds investigated here **(Figure 3)**. Rifampicin induced the protein and mRNA levels of CYP2C9 by 1.4- and 2.4-fold **(Figure 3)**. Previous reports employing 2D monolayer PHHs have found typically low induction, ~2-fold, of *CYP2C9* mRNA by rifampicin.^8,10,35,36^ The variable and low dynamic range of *CYP2C* mRNA induction is a challenge for accurate prediction of DDIs, particularly when the threshold for positive induction is generally regarded as 2-fold over control.^3,4,8,9^ In human DDI studies, rifampicin increased the clearance of CYP2C9 probe drugs flurbiprofen, tolbutamide and S-warfarin by a maximum of 2-fold^37–39^, which aligns well with our results for rifampicin that induced CYP2C9 protein and mRNA by 1.4- and 2- fold, respectively **(Figure 3)**. The high baseline expression of CYP2C9 protein in spheroids **(Figure S3)** may affect the extent of CYP2C9 induction found at the protein level **(Figure 3B)**. Finally, we did not find induction of CYP2C19 at the mRNA or protein level in one of three donors by any compound **(Figure 3)**. Previous studies in 2D monolayer PHHs have also reported donors with non-inducible CYP2C19 after rifampicin treatment.^8–10^ *CYP2C19* genotypes may affect the extent of induction in humans^40^, which might explain why some individuals have non-inducible CYP2C19.

Rifampicin induced the formation of specific metabolites of the Basel cocktail drugs in humans by ~1.2-, ~2-, ~4-, and ~4-fold for CYP2D6, CYP2B6, CYP2C19, and CYP3A4, respectively.^41^ We found similar induction here **(Figure 6)** for the enzyme activities of CYP2B6, CYP2C19 and CYP3A4, while CYP2D6 (formation of α-hydroxymetoprolol) was induced by 2-fold. Induction of CYP3A4-mediated formation of α-hydroxymetoprolol may play a role in 3D spheroid PHHs^42^, since rifampicin did not induce the mRNA or protein of CYP2D6 in any donor **(Figure 3)**. Induction of enzyme activities also underlines the translational potential of 3D spheroid PHHs.

Clinical relevance of transporter induction is not fully established, particularly for hepatic transporters.^5,43^ Studies have found ~2-fold upregulation of P-gp and MRP2 in the human liver after rifampicin administration.^43^ Here, rifampicin induced the mRNA of *ABCB1* and *ABCC2* by ~2-fold in all three donors **(Figure 4A)**, which agrees with the clinical findings. Moreover, our results are in line with two previous studies employing sandwich cultured PHHs and evaluating the inducing effects of rifampicin, phenobarbital and omeprazole on the expression of uptake and efflux transporters.^44,45^ We also found downregulation of *SLC22A1*, *SLC22A7* and *SLCO1B3* by omeprazole **(Figure 4A)** similar to the previous studies in sandwich cultured PHHs^44,45^, although the downregulation did not translate to the protein levels of OCT1 **(Figure 4B)**.

The main limitation of 3D spheroid PHHs is that not all batches of PHHs are able to form spheroids.^21^ Here, we utilized PHHs that were pre-qualified for formation of 3D cultures, however, comprehensive characterization of new batch of PHHs is a prerequisite for a viable 3D spheroid culture.^21^ Several other advanced hepatic culture systems have been developed within the past decade to allow long-term and stable culture of PHHs. Such models include microphysiological culture systems, different liver-on-chips and micropatterned co-cultures of hepatocytes and fibroblasts.^13,46^ A study compared the induction response of CYP2C8, CYP2C9 and CYP3A4 between 2D monolayer PHHs and advanced, long-term co-culture of fibroblasts and hepatocytes employing the same hepatocyte lots.^46^ The differences in maximal induction responses were within two-fold between the culture formats. Thus, the quality of batch of PHHs may be an independent factor that affects the induction responses regardless of culture format. In other words, we cannot exclude that the pre-qualification and characterization of PHH lots for 3D spheroid culture itself independently affects the more physiologically relevant magnitudes of induction found here than typically observed for 2D monolayer PHHs.

The main advantage of the 3D spheroid PHHs is the simple culture format in 96- or 384-well plates without additional devices and materials typically needed for other advanced hepatic cultures. The well plate format allows high throughput and scalability of assays. However, the low number of cells per spheroid, typically 1,500 to 5,000 hepatocytes, limits detection of low-clearance compounds, which was found here for the metabolites of caffeine and losartan. Some advancements have been made within the plate formats and multi-spheroid wells could improve the detection of slowly formed metabolites.^26^

In conclusion, we comprehensively investigated induction of eight important drug metabolizing CYPs and several efflux and uptake transporters in 3D spheroid PHHs by eight different inducers. We measured the relative mRNA and protein induction and found reproducible induction responses between three donors. Overall, we show excellent agreement between our *in vitro* and clinical DDI studies, which highlights the translational promise of this model.

## STUDY HIGHLIGHTS

What is the current knowledge on the topic?

- 2D monolayer cultured primary human hepatocytes (PHHs) have been the gold standard for *in vitro* induction studies but they rapidly dedifferentiate and lose expression of cytochrome P450 enzymes (CYPs). In contrast, 3D spheroid PHHs have an *in vivo* like phenotype and as an emerging model for *in vitro* studies of hepatic drug metabolism they may improve *in vitro* to *in vivo* translation.

What question did this study address?

- We investigated the 3D PHH spheroid model as an instrument for prediction of drug induced expression of CYPs and transporters.

What does this study add to our knowledge?

- We conclude that 3D spheroid PHHs are a suitable and versatile tool to investigate induction of CYPs and transporters *in vitro*. CYP3A4 induction responses by several different compounds were within clinically relevant magnitudes for the induction of mRNA, protein and enzyme activity.

How might this change clinical pharmacology or translational science?

- 3D spheroid PHHs may produce more physiological relevant results of drug-mediated induction and aid better prediction of drug-drug interactions in humans thus closing the translational gap.

## ACKNOWLEDGEMENTS

Rasmus Andersen is acknowledged for his excellent help during sample preparations.

## SUPPLEMENTARY INFORMATION

Supplementary Figures S1-S11, Tables S1-S4 and methods.

## AUTHOR CONTRIBUTIONS

EJ and TBS wrote manuscript, EJ and TBS designed research, EJ and HH performed research, EJ and HH analyzed data, OP and MIS contributed new reagents/analytical tools

