## Supplementary for "3D Spheroid Primary Human Hepatocytes for Prediction of Cytochrome P450 and Drug Transporter Induction"

### SUPPLEMENTARY FIGURES

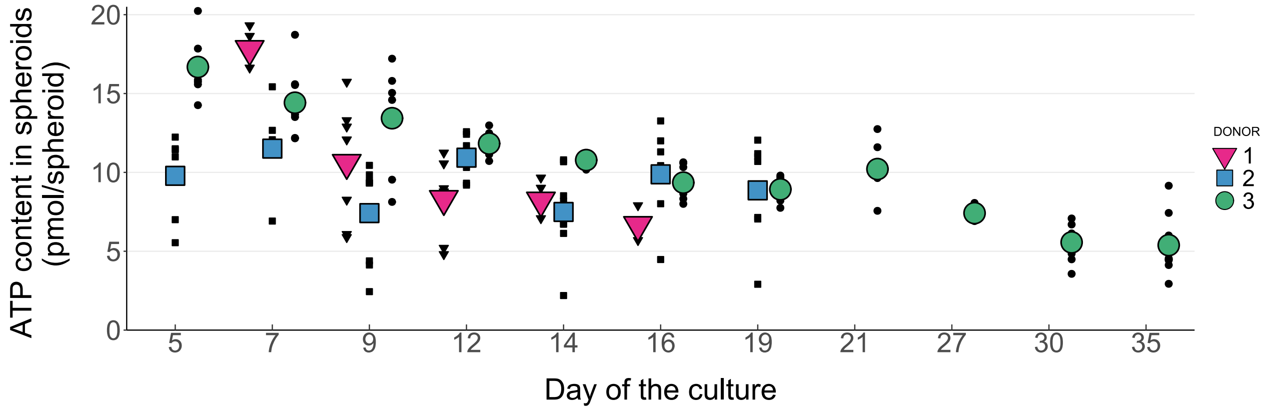


**FIGURE S1**. *Viability of spheroids from day 5 to 35*. ATP content of spheroids was determined for each donor on days 5, 7, 9, 12, 14 and 16 after seeding hepatocytes on ultra-low attachment plates. Data for donor 1 on day 5 and 19, and data for donor 1-2 on days 21-35 are missing. Data for days 5-16 are same as in Figure 2. N = 4-8 biological replicates (spheroids) per donor and time-point.

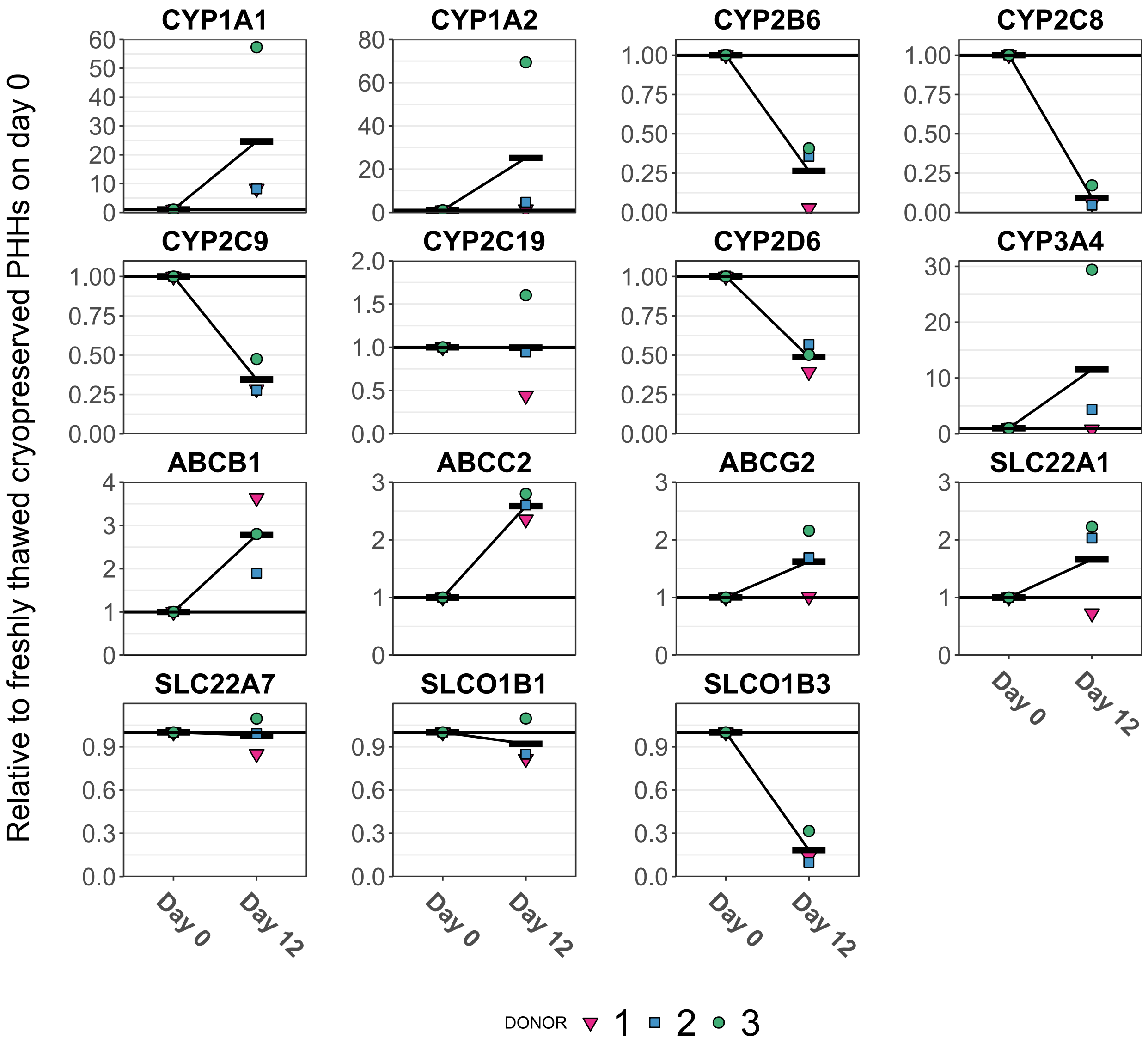


**FIGURE S2**. *Relative mRNA expression of CYPs and transporters in thawed hepatocytes versus the culture day 12 of spheroids*. Expression of mRNA levels were measured from freshly thawed cryopreserved PHHs (Day 0) or from spheroids on day 12 (Day 12). For each hepatocyte donor, the expression values of individual genes and proteins were normalized to the mean of Day 0 values. N = 3 hepatocyte donors presented as a mean value of three biological replicates of 24 000 cells or pools of 16 spheroids. Horizontal lines present mean of each group.

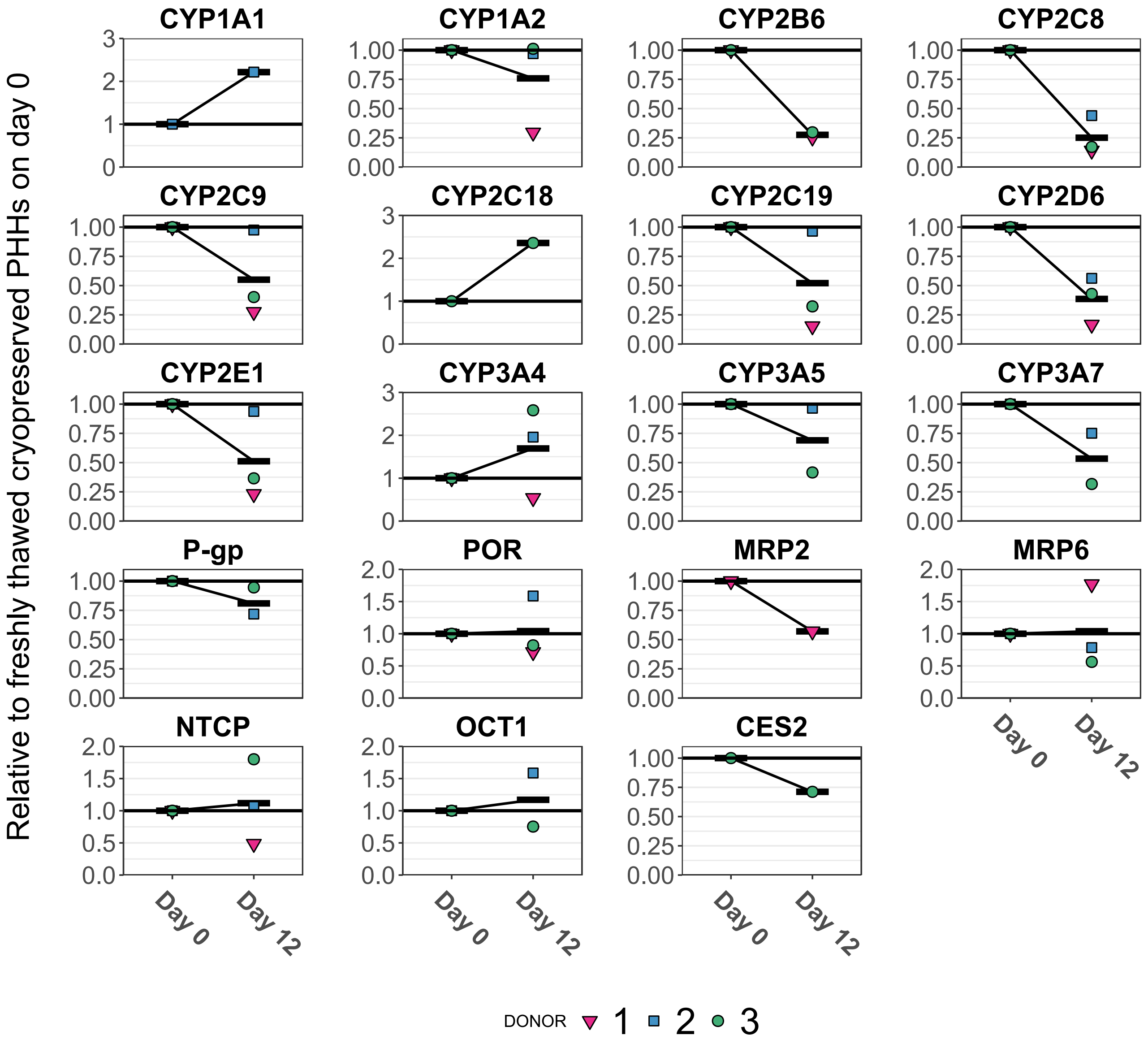


**FIGURE S3**. *Relative protein expression of CYPs and transporters in thawed hepatocytes versus the culture day 12 of spheroids*. Expression of protein levels were measured from freshly thawed cryopreserved PHHs (Day 0) or from spheroids on day 12 (Day 12). Some protein analysis samples were below limit of quantification, such as CYP2B6 in donor 2. For each hepatocyte donor, the expression values of individual genes and proteins were normalized to the mean of Day 0 values. N = 3 hepatocyte donors presented as a mean value of three biological replicates of 24 000 cells or pools of 16 spheroids. Horizontal lines present mean of each group.

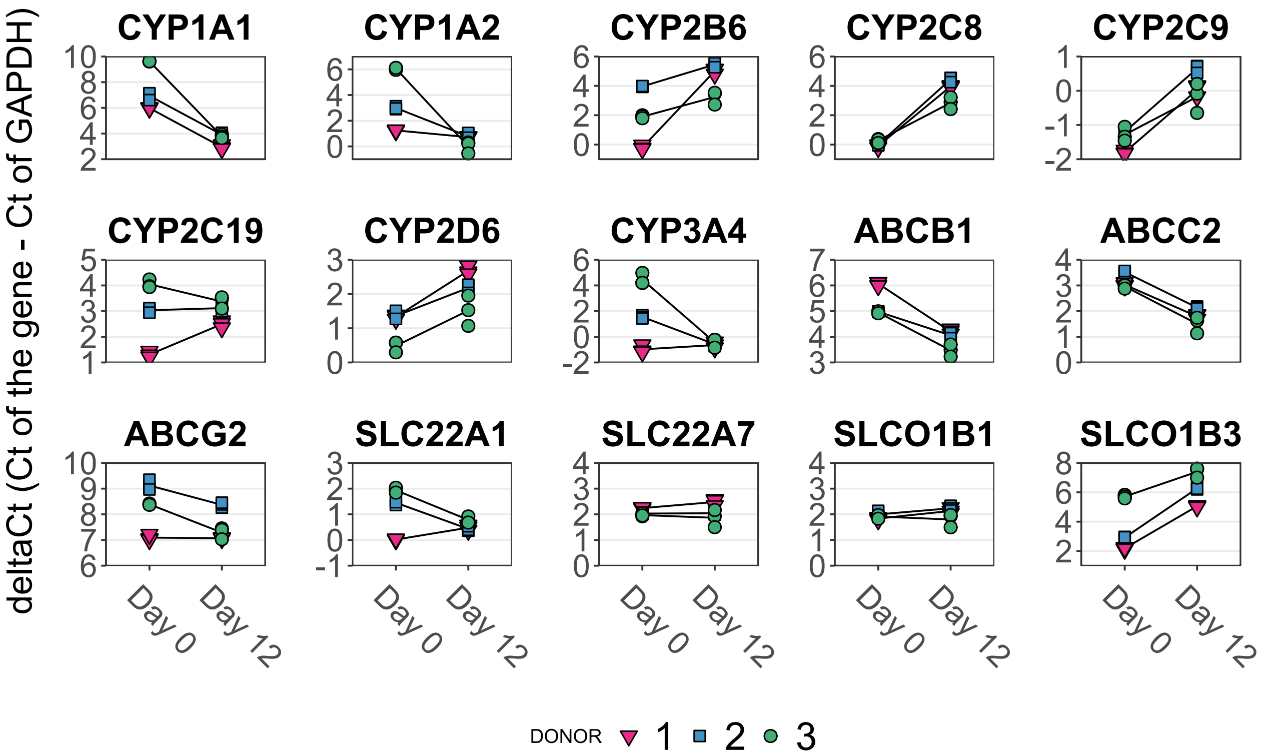


**FIGURE S4**. *Semi-quantitative analysis of CYP and transporter mRNAs in freshly thawed cryopreserved PHHs on day 0 and 3D spheroid PHHs on day 12*. Expression of mRNA was measured from cryopreserved hepatocytes after thawing (Day 0) or from spheroids on 12 day (Day 12). Raw Ct-values were normalized by the expression of GAPDH resulting in deltaCt values. **Note that $2^{-deltaCt}$ transformation is needed to convert deltaCt values to a linear scale.** N = 3 biological replicates each including 24 000 cells or 16 spheroids per sample and donor. Lines connect the means between day 0 and day 12 for each donor.

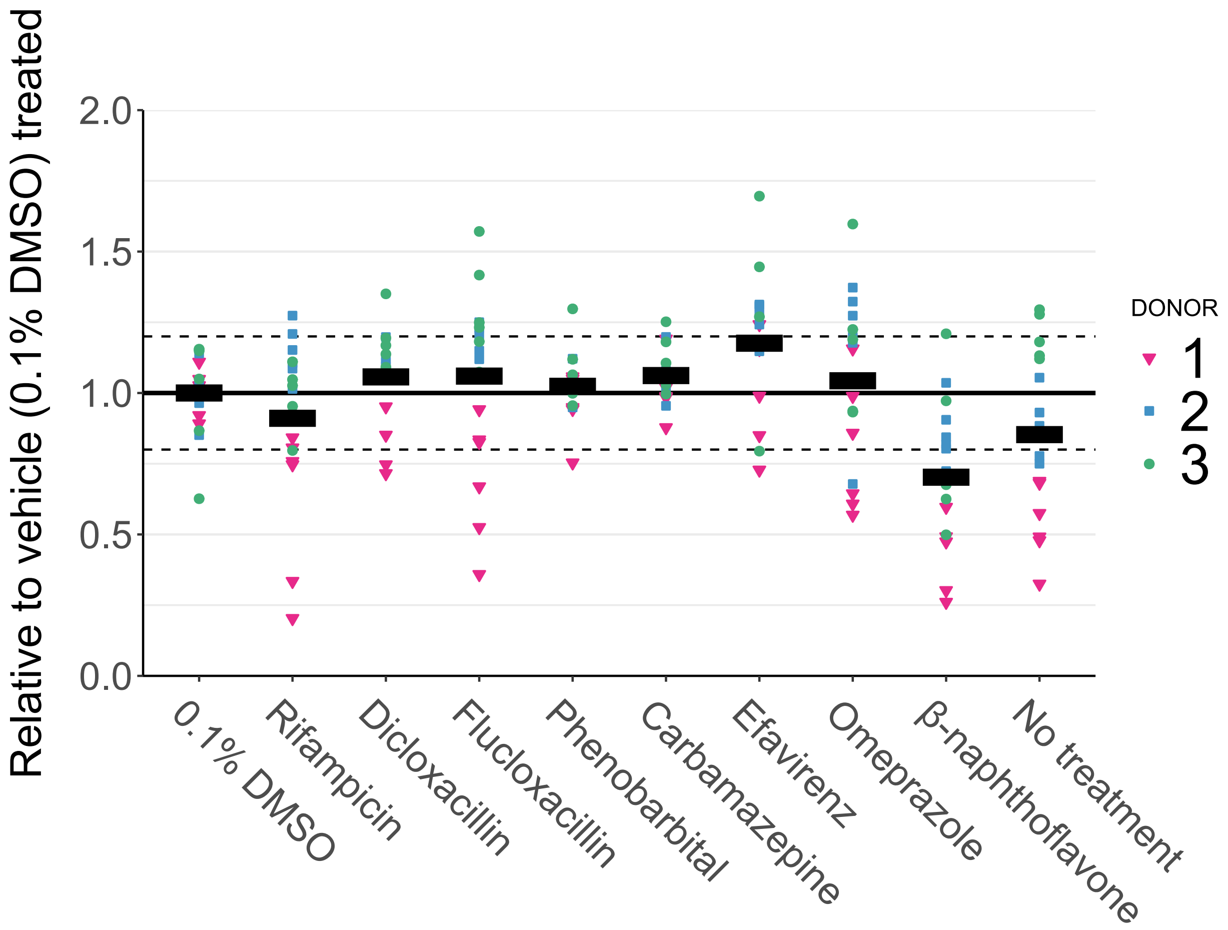


**FIGURE S5**. *Effect of induction treatments on spheroid viability.* ATP content of spheroids was measured for each donor after 4 days of treatment on day 12 and normalized to the mean of vehicle control (0.1% DMSO). Dashed lines indicate 20% intervals relative to the vehicle control. Data are summarized as means. N = 4-6 biological replicates per donor and treatment.

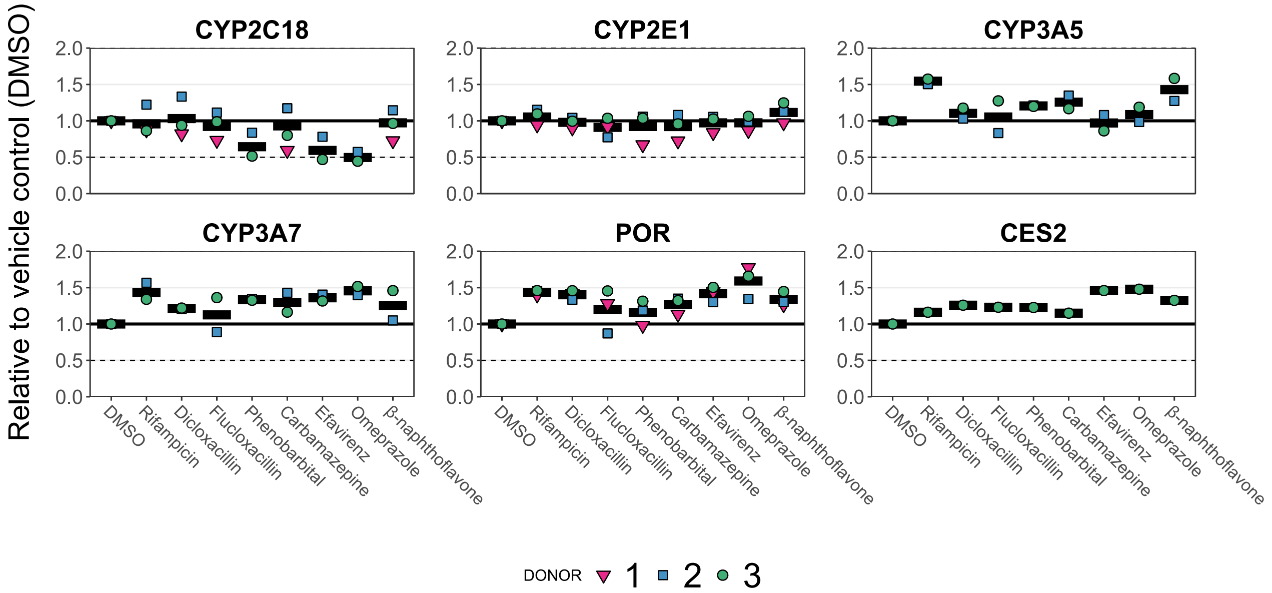


**FIGURE S6**. *Compound-specific induction of additional hepatic CYPs and CES2 at protein levels in 3D spheroid PHHs*. Induction of protein levels were determined after four days of treatment with the compounds. For each donor, induction was normalized to the mean response of 0.1% DMSO group. Data are summarized as means of three hepatocyte donors. Dashed lines present two-fold difference to the DMSO group. Some protein samples were below limit of quantification, such as CYP3A5 in donor 1, or were not analyzed, such as CES2 in donors 1 and 2. N = 3 hepatocyte donors each presented as a mean value of three biological replicates of pools of 16 spheroids.

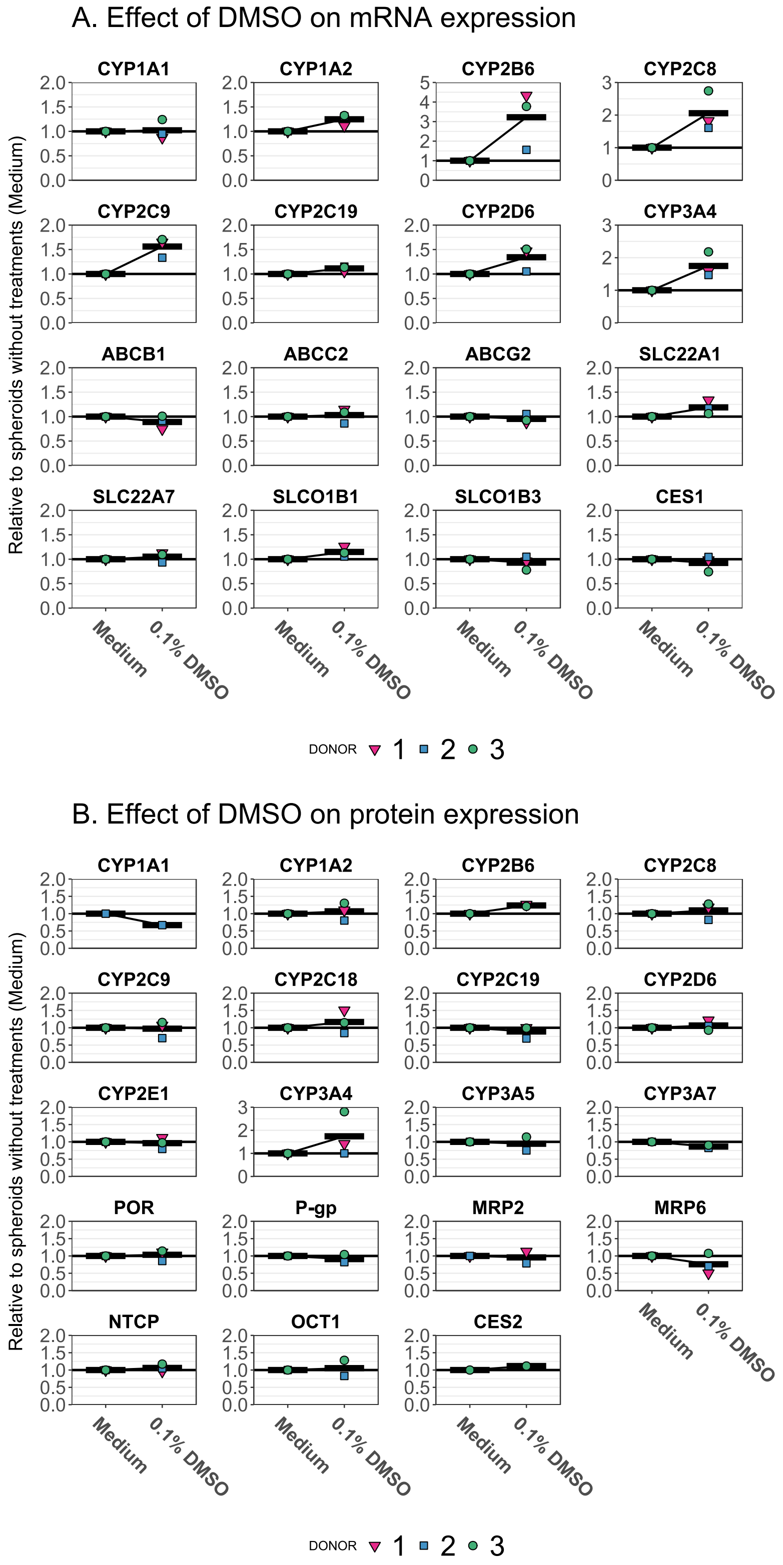


**FIGURE S7**. *Effect of 0.1% DMSO treatment on the expression of CYPs and drug transporters analyzed*. Expression of mRNA (A) or protein (B) were measured from spheroids without any treatment (Medium) or spheroids treated with vehicle for induction studies (0.1% DMSO). For each donor, the values were normalized to mean of medium group. Data are summarized as means of three hepatocyte donors and lines connect the means between groups. N = 3 hepatocyte donors each presented as a mean value of three biological replicates of pools of 16 spheroids. Data for 0.1% DMSO groups are same as presented in Figures 3, 4 and S6, and data for Medium groups are same as presented in Figures S2 and S3 (Day 12). For protein analysis, some samples were below limit of quantification, for example CYP3A5 in donor 2, and are missing from figures in panel B.

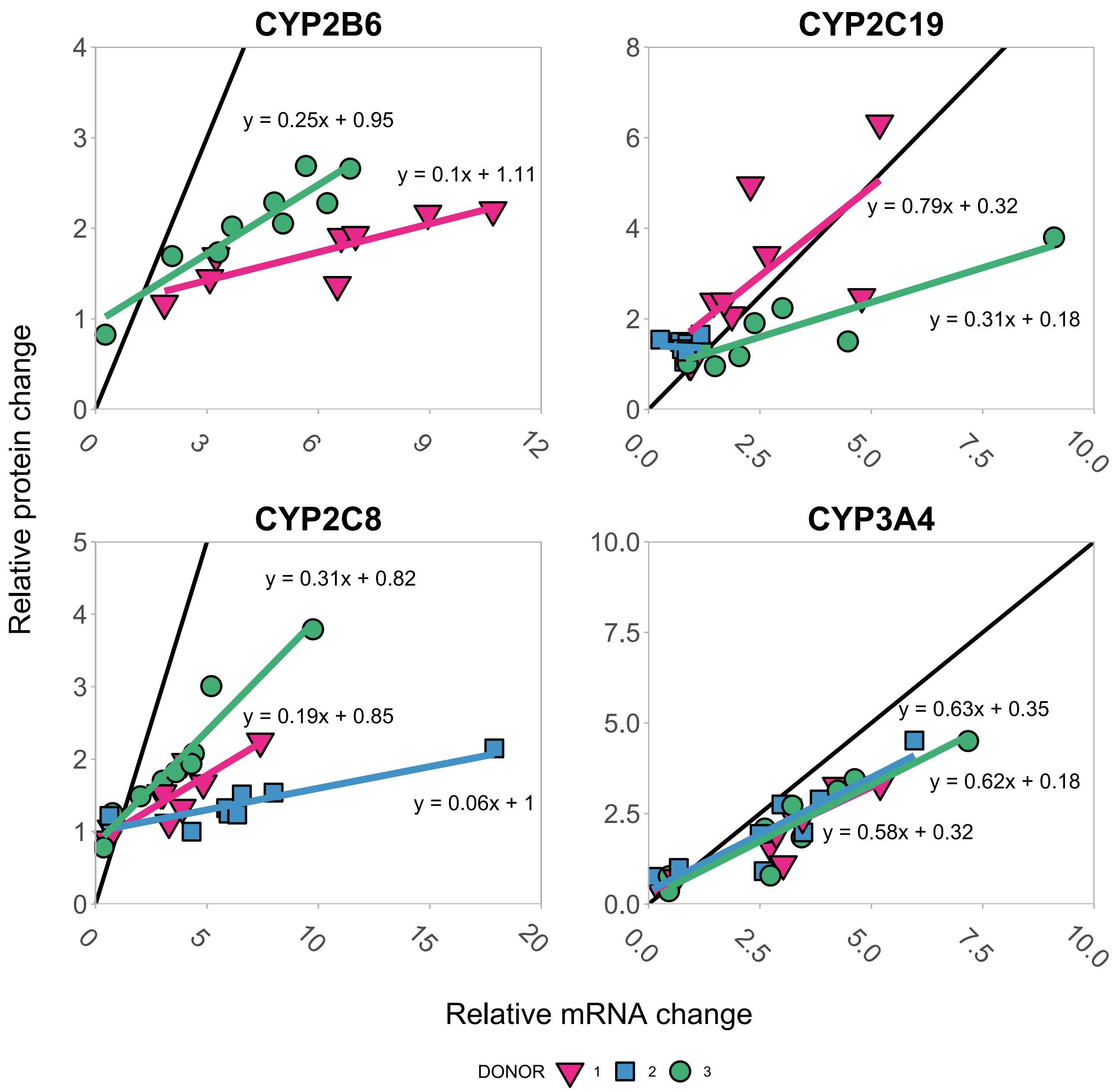


**FIGURE S8**. *Regression analysis between the relative mRNA and protein induction*. Analysis was conducted for CYP2B6, CYP2C19, CYP2C8 and CYP3A4. Each data point presents the mean of relative induction at mRNA (x axis) and protein (y axis) level for one of the compounds, excluding 0.1% DMSO, from Figure 3. Colored lines represent a linear regression (protein change = k * mRNA change + b) for each donor separately. Black lines represent the line of unity.

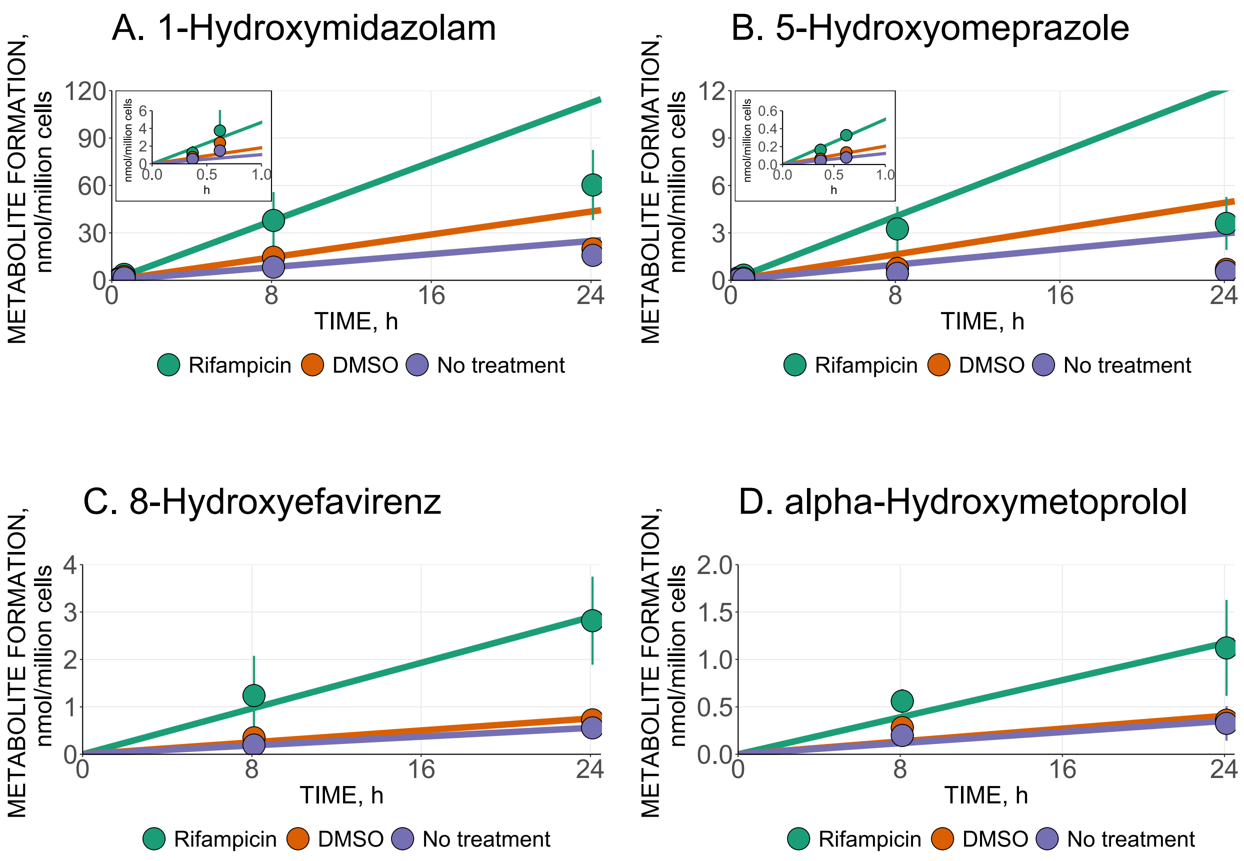


**FIGURE S9**. *Time-linearity of CYP3A4, CYP2C19, CYP2B6 and CYP2D6 enzyme activity in the Basel cocktail incubations*. Spheroids were treated with 10 µM rifampicin (Rifampicin), vehicle (0.1% DMSO) or without treatment (No treatment) for 4 days before incubation with the Basel cocktail (10 µM midazolam, 30 µM omeprazole, 20 µM efavirenz, 40 µM metoprolol, 30 µM losartan and 160 µM caffeine) for ~0.25, ~0.5, 8 and 24 hours. Formation of specific metabolites by CYP3A4 (A, 1-hydroxymidazolam), CYP2C19 (B, 5-hydroxyomeprazole), CYP2B6 (C, 8-hydroxyefavirenz) and CYP2D6 (D, α-hydroxymetoprolol) were measured from samples. A linear fitting (metabolite formation = k * time) was conducted for 0.37, 0.62 and 8.1 hours (A), 0.37 and 0.62 hours (B) or 8.1 and 24.1 hours (C and D). Data are presented as mean ± SD. N = 2 biological replicates both including 3 spheroids per condition.

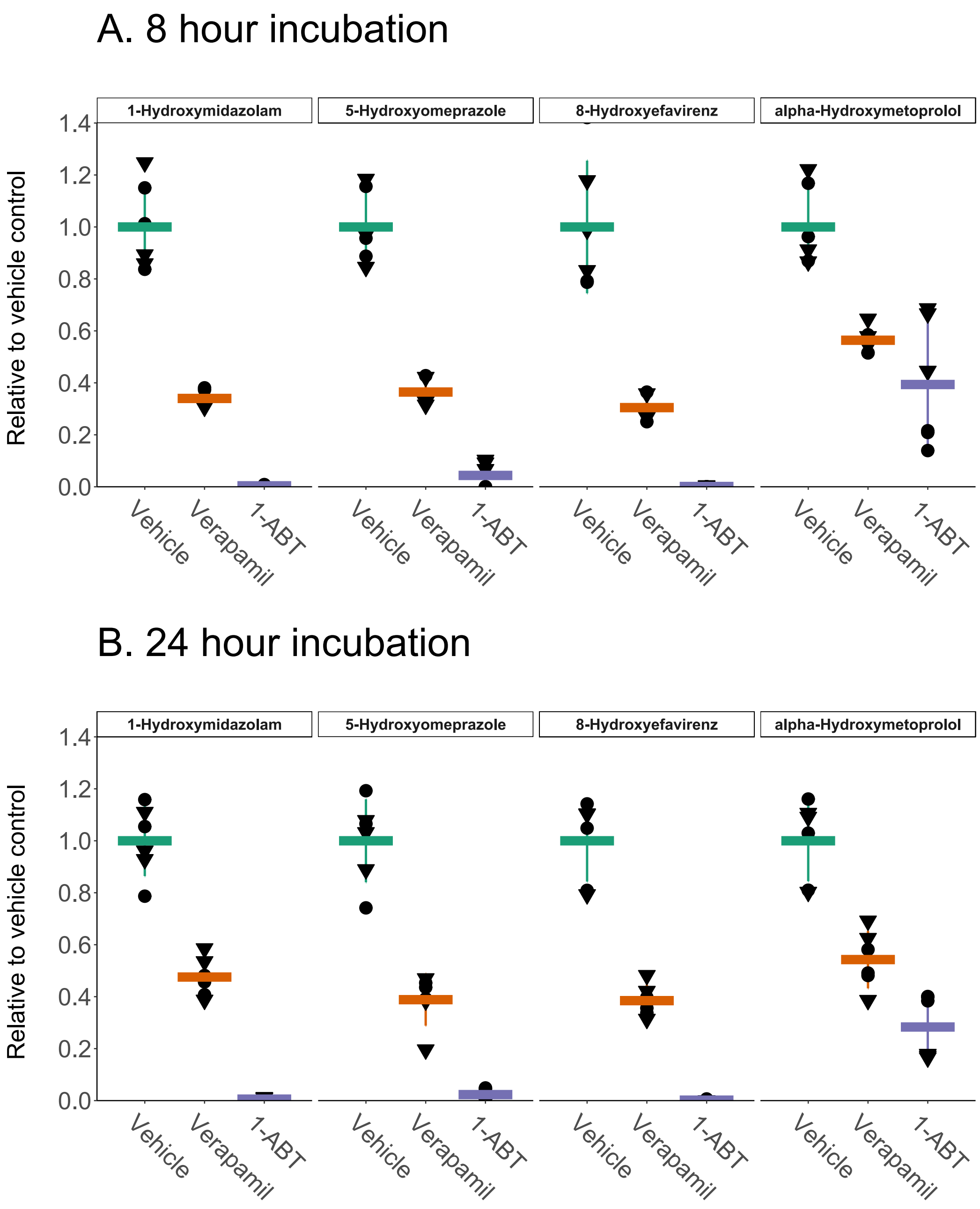


**FIGURE S10**. *Inhibition of CYP3A4, CYP2C19, CYP2B6 and CYP2D6 activities in the Basel cocktail incubations*. Spheroids were pre-incubated with 0.1% DMSO (Vehicle), 25 µM verapamil (Verapamil) or 1000 µM 1-aminobenztriazole (1-ABT) for 2 to 3 hours before applying the Basel cocktail (10 µM midazolam, 30 µM omeprazole, 20 µM efavirenz, 40 µM metoprolol, 30 µM losartan and 160 µM caffeine) to spheroids, together with the inhibitors or vehicle, for further incubation of 8 (A) or 24 hours (B). The final DMSO concentration was 0.15%. Data were normalized to mean MS-responses of vehicle groups and are summarized as mean ± SD. N = 2 biological replicates both including 3 spheroids per condition. Triangles and points represent donor 1 and 3.

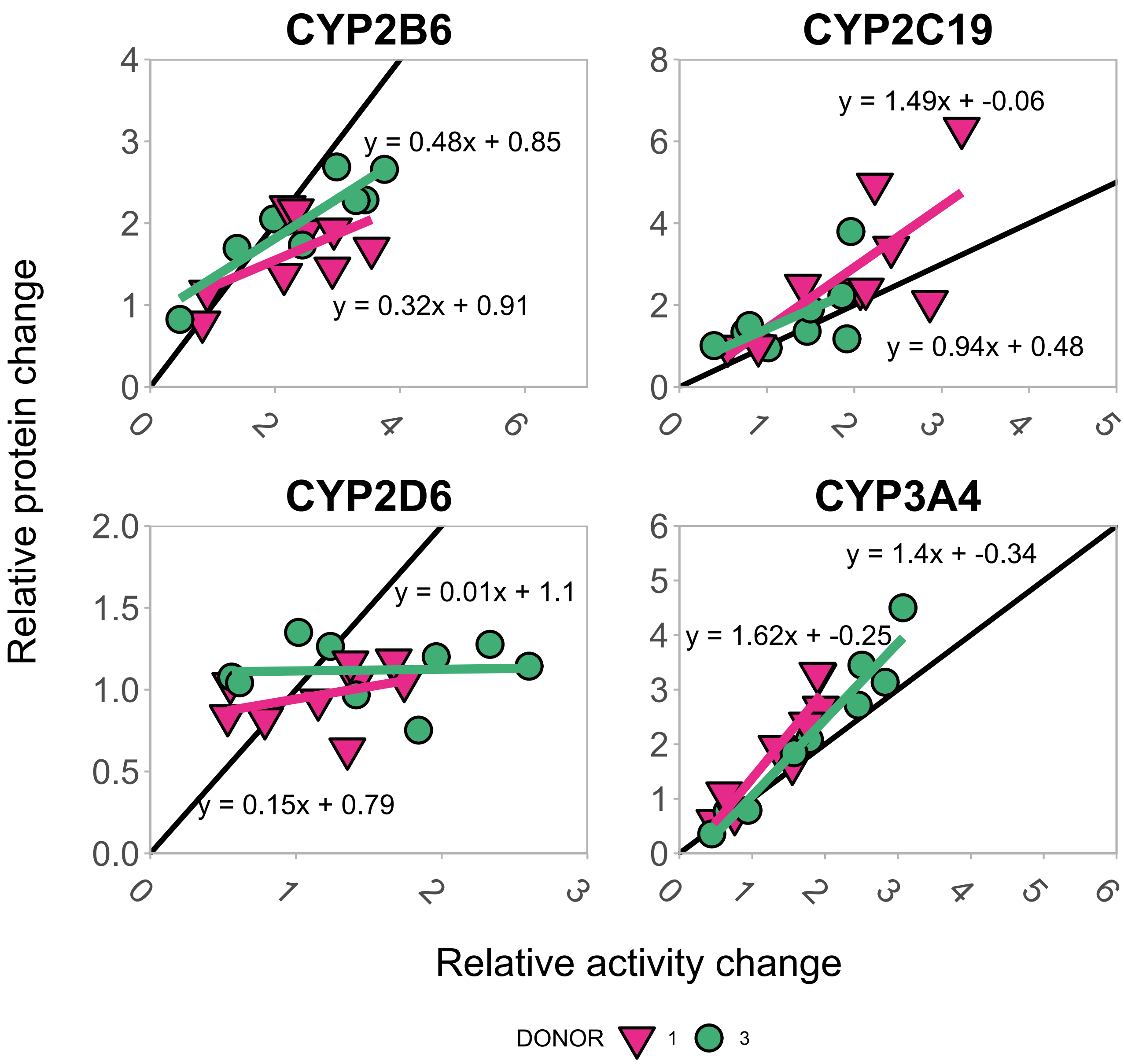


**FIGURE S11**. *Regression analysis between the relative induction of enzyme activity and protein levels*. Analysis was conducted for CYP2B6 (A), CYP2C19 (B), CYP2D6 (C) and CYP3A4. Each data point presents the mean of relative induction at enzyme activity (x axis) and protein (y axis) level for one of the compounds, excluding 0.1% DMSO, from Figures 3B and 6. Colored lines represent a linear regression (protein change = k * enzyme activity change + b) for each donor separately. Black lines represent the line of unity.

### SUPPLEMENTARY TABLES

**Table S1**. Characteristics of hepatocyte lots employed in this study. The information is from data sheets provided by the supplier of PHHs (Thermo Fisher Scientific, Walthan, MA, USA).

| **Donor number in this manuscript** | **LOT** | **Gender** | **Race** | **Age** |
|---|---|---|---|---|
| 1 | HU8345-A | Male | Asian | 47 |
| 2 | HU8339-A | Female | African American | 31 |
| 3 | HU8373-A | Female | Caucasian | 26 |

**Table S2**. LC-MS analysis of the specific metabolites from spheroid samples incubated with the Basel cocktail. Exact masses, retention times and commercial sources of the metabolites and their internal standards (labelled compounds) are included. 8-hydroxyefavirenz and its internal standard were analyzed in negative ionization mode, while positive ionization mode was employed for all the other compounds.

| **Metabolite/internal standard** | **Exact mass, m/z** | **Retention time, min** | **Source** |
|---|---|---|---|
| Paraxanthine | 181.0720 | 2.2 | Sigma-Aldrich (St. Louis, MO, USA) |
| Paraxanthine-$d_6$ | 187.1097 | 2.1 | Sigma-Aldrich |
| α-hydroxymetoprolol | 284.1856 | 2.9 | Sigma-Aldrich |
| Metoprolol-$d_7$ | 275.2347 | 4.7 | Sigma-Aldrich |
| 5-hydroxyomeprazole | 362.1169 | 5.2 | Toronto Research Chemicals (Toronto, ON, Canada) |
| Omeprazole-$d_3$ | 349.1408 | 5.7 | Toronto Research Chemicals |
| 1-hydroxymidazolam | 342.0804 | 6.3 | Chiron AS (Trondheim, Norway) |
| Midazolam-$d_7$ | 333.1294 | 6.1 | Chiron AS |
| Losartan carboxylic acid | 437.1487 | 7.6 | Toronto Research Chemicals |
| Losartan-$d_4$ carboxylic acid | 440.1676 | 7.6 | Toronto Research Chemicals |
| 8-hydroxyefavirenz | 330.0150 | 7.9 | Toronto Research Chemicals |
| 8-hydroxyefavirenz-$d_4$ | 334.0401 | 7.9 | Toronto Research Chemicals |

**Table S3**. Surrogate peptides, UniProt ID and LC-gradients for the proteins quantified in this study.

| **Protein** | **Surrogate peptide** | **UniProt ID** | **LC-gradient#** |
|---|---|---|---|
| ABCB1 | EANIHAFIESLPNK | P08183 | 10 min |
| ABCC2 | LTIIPQDPILFSGSLR | Q92887 | 10 min |
| ABCC6 | ISIIPQDPILFPGSLR | O95255 | 10 min |
| CES2 | APVYFYEFQHQPSWLK | O00748 | 10 min |
| POR | ESSFVEK | P16435 | 10 and 20 min |
| CYP1A1 | GFYIPK | P04798 | 10 and 20 min |
| CYP1A2 | DTTLNGFYIPK | P05177 | 10 and 20 min |
| CYP2B6 | AEAFSGR | P20813 | 10 and 20 min |
| CYP2C18 | EALIDHGEEFSGR | P33260 | 10 and 20 min |
| CYP2C19 | GHFPLAER | P33261 | 10 and 20 min |
| CYP2C8 | EALIDNGEEFSGR | P10632 | 10 and 20 min |
| CYP2C9 | GIFPLAER | P11712 | 10 and 20 min |
| CYP2D6 | GTTLITNLSSVLK | P10635 | 10 and 20 min |
| CYP2E1 | DEFSGR | P05181 | 10 and 20 min |
| CYP3A4* | LQEEIDAVLPNK* | P08684 | 10 and 20 min |
| CYP3A5 | EIDAVLPNK | Q9HB55 | 10 and 20 min |
| CYP3A7 | EIDTVLPNK | P24462 | 10 and 20 min |
| SLC10A1 | GIYDGDLK | Q14973 | 12 min |
| SLC22A1 | LSPSFADLFR | O15245 | 10 min |

#10 and 20 min gradients were previously described[1], and 10 min gradient is an elongated version of a 6 min method previously described for Acclaim Pepmap RSLC C18 column[2]
*Sequence is also present in CYP3A43. No or very low expression of CYP3A43 has been observed in the liver.

**TABLE S4**. *mRNA and protein expression of CYPs and transporters in advanced culture formats of PHHs normalized to freshly thawed hepatocytes on day 0.* A mean value of expressions was calculated for this study and each literature reference. The red and green values indicate expressions below or above 2-fold difference in comparison to freshly thawed hepatocytes.

| | **This study on day 12.** | | Spheroids on day 14.[3] | Spheroids on day 14.[4] | Spheroids on day 14.[5] | | Spheroids (hepatocytes + non-parenchymal cells) in a propietary medium on day 14.[6] | | Micropatterned co-cultures of hepatocytes and mouse fibroplasts on day 7 (Hepatopac).[7] | Liver on chip on day 7.[8] | Liver on chip on day 6.[9] |
|---|---|---|---|---|---|---|---|---|---|---|---|
| **CYP/ Transporter** | **mRNA** | **Protein** | **mRNA** | **Protein** | **mRNA** | **Protein** | **mRNA** | **Protein** | **mRNA** | **mRNA** | **mRNA** |
| CYP1A1 | 24.6 | 2.2 | | 0.26 | | | 19 | | | | 1.78 |
| CYP1A2 | 25.2 | 0.76 | 114 | 0.62 | 1.07 | 1.46 | 9 | 0.31 | 0.31 | 0.71 | 0.37 |
| CYP2B6 | 0.26 | 0.28 | | 0.3 | 0.19 | 1.28 | 1.9 | 0.6 | 0.09 | 1.3 | 0.09 |
| CYP2C8 | 0.09 | 0.25 | 0.4 | 0.51 | 0.11 | 1.36 | 1 | 0.2 | 0.16 | 0.27 | 0.13 |
| CYP2C9 | 0.35 | 0.55 | 0.77 | 0.56 | 0.44 | 1.5 | 1.3 | 0.22 | 0.4 | 0.27 | 0.26 |
| CYP2C19 | 1.0 | 0.52 | | 0.51 | 0.34 | 1.74 | 1.5 | | | 2.7 | 0.37 |
| CYP2D6 | 0.49 | 0.39 | 0.37 | 0.31 | 0.9 | 1.5 | 0.91 | 0.15 | 0.47 | 0.32 | 0.74 |
| CYP3A4 | 11.8 | 1.7 | 95 | 0.81 | 0.29 | 1.17 | 1.2 | 0.25 | 0.44 | 1.19 | 1.38 |
| ABCB1 | 2.8 | 0.81 | | 0.77 | | | 1.5 | 2.2 | 0.89 | 3.3 | 1.54 |
| ABCC2 | 2.6 | 0.57 | 4.6 | 1.15 | | | 1.7 | 2.7 | | 2.7 | |
| ABCG2 | 1.6 | | | | | | 1.2 | | | 0.74 | |
| SLC22A1 | 1.7 | 1.1 | | 0.75 | | | 0.79 | 0.16 | 0.62 | 0.59 | |
| SLC22A7 | 0.98 | | | 0.37 | | | | | | | |
| SLCO1B1 | 0.92 | | 2 | 0.59 | | | 1.8 | 0.54 | | 0.44 | |
| SLCO1B3 | 0.18 | | | 0.63 | | | 0.11 | | | 0.08 | |

### SUPPLEMENTARY METHODS

##### 3D SPHEROID CULTURE OF PRIMARY HUMAN HEPATOCYTES

A vial of PHHs was thawed on day 0 in a +37˚C water bath until a small clump of ice remained. Cell suspension was poured into 45 ml of hepatocyte thawing medium (CM7000 or CM7500) and centrifuged 10 min at 100 g. The supernatant was removed, and cells were resuspended in William's E medium without phenol red supplemented with primary hepatocyte thawing and plating supplements (CM3000). This plating medium contains 5% fetal bovine serum, 1 µM dexamethasone, 100 U/ml penicillin, 100 µg/ml streptomycin, 5 µg/ml human recombinant insulin, 2 mM GlutaMAX (L-glutamine) and 15 mM HEPES. Cell concentration and viability was determined by trypan blue staining and manual counting with a hemocytometer. The viability of PHHs was between 91 and 93%. Cells were diluted in the plating medium and 1,500 viable cells in a volume of 100 µl was transferred to each well of a 96-well ultra-low attachment plate (Nunclon Sphera). The plates were centrifuged 2 min at 200 g and then transferred to a cell culture incubator (5% $CO_2$ and +37˚C) for five days.

PHHs aggregated to spheroids within five days and on days 5 to 7 the plating medium was changed to a maintenance medium by removing 50 µl or 70 µl (day 6 and 7) and adding 70 µl of new medium. The maintenance medium (donor 3) contains 0.1 µM dexamethasone, 100 U/ml penicillin, 100 µg/ml streptomycin, 10 µg/ml human recombinant insulin, 5.5 µg/ml transferrin, 6.7 ng/ml selenium and 2 mM GlutaMAX (L-glutamine). In the case of donors 1 and 2, the maintenance medium was recommended in the protocol of 3D spheroid PHHs by Thermo Fisher Scientific and consisted of William's E supplemented with primary hepatocyte maintenance supplements (CM4000) resulting in 50 U/ml penicillin, 50 µg/ml streptomycin, 6.25 µg/ml human recombinant insulin, 6.25 µg/ml transferrin, 6.25 ng/ml selenium, 1.25 mg/ml bovine serum albumin, 5.35 µg/ml linoleic acid and 15 mM HEPES, while the other components were same. This supplement is no longer recommended for 3D spheroid PHHs (communication with ThermoFisher Scientific). After day 7, 70% of the medium was changed every second or third day.

#### CYP ENZYME ACTIVITY AND LIQUID CHROMATOGRAPHY-MASS SPECTROMETRY ANALYSIS OF THE METABOLITES

Metabolites from the enzyme activity samples were analyzed with liquid chromatography-mass spectrometry analysis (LC-MS). First, the samples were subjected to three freeze-thaw cycles and vortexing to break spheroids. 50 µl of samples were de-conjugated with 62.5 units of β-Glucuronidase from Escherichia coli Type IX-A (Sigma-Aldrich) by incubating them overnight at +37˚C. After incubation, proteins were precipitated with 150 µl of ice-cold methanol containing internal standards (paraxanthine-$d_6$, 8-hydroxy-efavirenz-$d_4$, losartan-$d_3$ carboxylic acid, omeprazole-$d_3$, metoprolol-$d_7$ and midazolam-$d_7$) and by incubating samples at -20˚C for 60 min or longer. Samples were centrifuged 10 min at 21 000 g and the resulting supernatants were diluted 1:1 with water and transferred to glass vials for LC-MS analysis.

The LC-MS system consisted of Vanquish LC system equipped with ACE Excel 3 C18-AR column and a pre-column filter (Advanced Chromatography Technologies, Aberdeen, UK) connected to Q Exactive Orbitrap mass spectrometer with a heated electrospray ionization source (ThermoFisher Scientific). The column and sample chambers were kept at +40˚C and +10˚C, respectively. Water and methanol, both containing 0.1% formic acid, were employed as chromatography eluents A and B with a flow rate of 0.4 ml/min. Injection volume of samples was 5 µl. The chromatography method was 10% B from 0 to 1 min and the analytes were eluted with a gradient from 10 to 95% B between 1 to 8.9 min, which was followed with a column wash and equilibrium step before a new injection. Ion-source parameters were set at 50 for S-lens RF level, +3 or -3 kV (8-hydroxy-efavirenz and its internal standard) for spray capillary voltage, 275˚C and 400˚C for capillary and auxiliary gas heater temperature, and 2, 11 and 46 for sweep, aux and sheath gas flow rates. The mass spectrometer was operated in targeted single ion monitoring multiplex mode with an inclusion list containing the m/z values of analytes. Orbitrap parameters were set at resolution of 70 000, AGC target of 5e5 and maximum ion time of 200 ms. Exact masses extracted for metabolites and internal standards, their retention times and commercial sources for standards are presented in Table S2. Standard curves for the metabolites were prepared by spiking increasing concentrations of metabolite standards into a matrix (the maintenance medium containing the Basel cocktail) and preparing them similar to the samples. Lower limit of quantification for each analyte was determined at a level producing at least a five-fold higher response than a blank sample containing only the matrix.

**RNA EXTRACTION AND qPCR**

The following TaqMan assays (Thermo Fisher Scientific) were employed in qPCR analysis: Hs02758991_g1 (*GAPDH*), Hs00153120_m1 (*CYP1A1*), Hs00167927_m1 (*CYP1A2*), Hs04183483_g1 (*CYP2B6*), Hs00946140_g1 (*CYP2C8*), Hs04260376_m1 (*CYP2C9*), Hs00426380_m1 (*CYP2C19*), Hs00164385_m1 (*CYP2D6*), Hs00604506_m1 (*CYP3A4*), Hs00184500_m1 (*ABCB1*), Hs00166123_m1 (*ABCC2*), Hs01053790_m1 (*ABCG2*), Hs00427552_m1 (*SLC22A1*), Hs00198527_m1 (*SLC22A7*), Hs00272374_m1 (*SLCO1B1*), Hs00251986_m1 (*SLCO1B3*) and -Hs00275607_m1 (*CES1*).

**INHIBITION STUDIES OF CYP ENZYME ACTIVITY IN 3D PHH SPHEROIDS**

Inhibitors were selected based on literature reports. 1-ABT completely abolishes the CYP3A4 activity in human hepatocytes.[10] In human liver microsomes, 1-ABT inhibits CYP2C19 and CYP2B6 and weakly CYP2D6.[11] Verapamil is a CYP3A4 mechanism-based inhibitor.[12] Verapamil also inhibits CYP2C19 and CYP2D6 in human liver microsomes.[13]

Studies on inhibition of CYP enzyme activities in 3D PHH spheroids were conducted essentially similarly as other activity assays. Verapamil and 1-aminobenztriazole were acquired from Sigma-Aldrich (St. Louis, MO, USA), and dissolved in DMSO as 1,000x concentrations. After two washes, the inhibitors and vehicle (DMSO) were applied to spheroids for a pre-incubation period of 2-3 hours. The activity assay was initiated by changing the medium to new media containing the Basel cocktail and 25 µM verapamil or 1000 µM 1-ABT or 0.1% DMSO. Final DMSO concentration in the incubation was 0.15% in each case. After 8- and 24-hour incubations, samples were collected and processed similarly to other enzyme activity samples. For data analysis, analyte to internal standard ratios for samples were extracted from LC-MS data and normalized by the mean value of this ratio in vehicle controls for each metabolite and time point.'

**LINEAR FITTINGS of DATA**

Linear fittings for the metabolite formation (Figure S9) were done with lm function in R (version 4.2.1, The R Foundation). The formula for the fittings was y ~ 0 + x, where y is metabolite formation and x is time. Collecting all samples for each time point lasted about seven minutes and for the linear fitting incubation timepoints were corrected for this time, e.g. 37 min was used instead of 30 min. The linear regression between the relative mRNA and protein induction (Figure S8), and the relative protein and enzyme activity induction (Figure S11) was done similarly with lm function in R. The formula was y ~ x, where y is the relative protein induction and x is the relative mRNA induction.
